# Epitope-Based Peptide Vaccine Design Against Fructose Bisphosphate Aldolase of Candida Glabrata: An Immunomics Approach

**DOI:** 10.1101/2020.07.03.180430

**Authors:** Elamin Elhasan LM, Mohamed B. Hassan, Reham M. Elhassan, Fatima A. Abdelrhman, Essam A. Salih, Asma Ibrahim. H, Amna A. Mohamed, Hozaifa S. Osman, Marwa Saad M. Khalil, Athar A. Alsafi, Abeer Babiker Idris, Mohamed A. Hassan

## Abstract

**Background:** *Candida glabrata* is a human opportunistic pathogen that can cause life-threatening systemic infections. Although, there are multiple effective vaccines against fungal infections, and some of these vaccines were engaged in different stages of clinical trials, none of them yet approved by (FDA).

**Aim:** To predict the most conserved and immunogenic B- and T-cell epitopes from the Fructose Bisphosphate aldolase (Fba1) protein of *C. glabrata*.

**Materials and Methods:** 13 *C. glabrata* Fructose bisphosphate aldolase protein sequences (361amino acid) were retrieved from NCBI and several in silico tools presented in the IEDB server for predicting peptides were used and homology modeling and molecular docking were performed.

**Result:** The promising B-cell Epitopes were AYFKPH, VDKESLYTK, and HVDKESLYTK. While, promising peptides which have the high affinity to MHC I binding were: AVHEALAPI, KYFKRMAAM, QTSNGGAAY, RMAAMNQWL and YFKEHGEPL. Two peptides (LFSSHMLDL and YIRSIAPAY) were noted to have the highest affinity to MHC class II that interact with 9 MHC class II alleles. The molecular Docking revealed the epitopes QTSNGGAAY and LFSSHMLDL have the high binding energy to MHC molecules

**Conclusion:** The epitope-based vaccines predicted by using immunoinformatics tools have remarkable advantages over the conventional vaccines that they are more specific, less time consuming, safe, less allergic and more antigenic. Further in vivo and in vitro experiments are needed to prove the effectiveness of the best candidates epitopes (QTSNGGAAY and LFSSHMLDL). To the best of our knowledge, this is the first study that has predicted B- and T-cells epitopes from Fba1 protein by using in silico tools in order to design an effective epitope-based vaccine against *C. galabrata*.

## 1. Introduction

Candidiasis is a fungal infection that has a high burden of morbidity and mortality in hospitalized and immunocompromised patients. It occurs in more than a quarter of a million patients every year with incidence rates for candidemia of 2–14 per 100000 ^[1–4]^. Candida pathogenicity is facilitated by a number of virulence factors, most importantly adherence to host surfaces including medical devices, biofilm formation and secretion of hydrolytic enzymes. Also, Candida cells elaborate polysaccharides, proteases, phospholipases and haemolysins that cause host cell damage which leads to the increase in the incidence and antifungal resistance of NCAC species, specifically *C. glabrata* and the unacceptably high morbidity and mortality associated with these species^[5]^.

*Candida glabrata* (*C. glabrata*) is a human opportunistic pathogen that can cause life-threatening systemic infections. *C. glabrata* is not polymorphic, grow as blastoconidia (yeast), lack of pseudohyphal formation so classified in the genus Torulopsis *C. glabrata* cells (1–4 μm in size)^[6]^ and forms glistening, smooth, and cream-colored colonies^[7]^. During infection, invade the innate immune system such as macrophages, which belong to the first line of defense against invading pathogens. *C.glabrata* able to modify its phagosomal compartment, avoiding full maturation, acidification and thus prevent the reaching of not the hostile phagolysosomal environment ^[8]^. *C. glabrata* is able to invade the bloodstream and different organs in a mouse model that has intragastrointestinal infections^[9]^. The genome of *C. glabrata* was published in 2004 by Dujon *et al*. 2004^[10]^. *C. glabrata* has a haploid genome that allows adaptation to a wide range of environments ^[5, 11, 12]^. Also, its genome contains more tandem repeats of genes than the other Nakaseomyces^[13]^ *C. glabrata* covers 67 genes encoding putative adhesin which is cell wall proteins, including the Epa family with 17 members^[11, 14]^. Most importantly examples are Epithelial Adhesin 1 (Epa1p)^[15]^ and Fructose bisphosphate aldolase protein (Fba1) which play an essential role in the pathogenicity of Candida species mainly in the adhesion of the pathogen to the host ^[16, 17]^.

Fba1 is a yeast cell wall protein that presents in multiple species of Candida^[18]^. e.g *C. glabrata, C. parapsilosis, C. tropicalis* and *C. albicans* fungal pathogen.^[19–22]^ It is a very important enzyme in the glycolytic pathway^[23–26]^ and multifunctional protein^[27]^ that can facilitate the attachment (adhesion) to human cells or abiotic surfaces^[28–30]^, protects Candida cells from the host’s immune system^[28]^ and promotes the detoxification of the ROS generated during the respiratory burst^[28, 29, 31]^. However, proteomics analysis revealed that Fba1 is the most abundant and stable enzyme in Candida. Moreover, it is considered one of the main immunodominant proteins^[32, 33]^ in Candida cells and tested in the murine model as protected protein against candida^[34]^ especially, *C. albicans*; also introduced immunity to *C. glabrata*^[35]^ therefore Fba1 is a potential antifungal target in yeast^[36]^. Multiple vaccines used Fba1 as immunogenic protein against different pathogens such as the lethal challenging *S. pneumonia, salmonella spp*., and *M. bovis*^[37–39]^.

The incidence of fungal infection has been increasing in the latest years, due to several factors such as misuse of broad-spectrum antibiotics, cytotoxic chemotherapy, immunocompromised patients, and transplantations^[40, 41]^. Invasive fungal infections are a major cause of global morbidity and mortality, accounting for about 1.4 million deaths per year ^[42]^. Systemic fungal infections cost the healthcare industry approximately $2.6 billion per year in the USA alone ^[43]^. However, Candida species pose a base problem in hospitals, according to Healthcare-Associated Infections (HAI) ^[44–47]^. Although, there are multiple effective vaccines against fungal infections, and some of these vaccines were engaged in different stages of clinical trials, none of them yet approved by (FDA) ^[48]^. Therefore, there is an urgent and crucial need to design vaccines against candida species that might improve the quality of life for immunosuppressed patients ^[49]^. The aim of this study is to predict the most conserved and immunogenic B- and T-cell epitopes from the Fba1 protein of *C. glabrata* by using in silico tools presented in the IEDB server. The Immunoinformatics approach has multiple benefits in comparison to other approaches by being affordability, safety, time-saving and clinically applicable using different computational software techniques ^[50–52]^. To the best of our knowledge, this is the first study that has predicted the best candidates of multiple epitopes of Fba1 protein against *C. glabrata*.

## 2. Materials and Methods

### 2.1 Retrieving of Fructose Bisphosphate Aldolase protein sequences

13 *Candida glabrata* Fructose bisphosphate aldolase protein sequences (361aa proteins) were retrieved from NCBI (https://www.ncbi.nlm.nih.gov/protein) Database on 21 January 2019 ^[17]^.

### 2.2 Determination of conserved regions

Multiple sequence alignment (MSA) was used to determine the conserved regions, the retrieved sequences were aligned by MSA using Clustal-W as Applied in the Bio-Edit ^[53]^.

### 2.3 Prediction of B-cell epitope

The reference sequence of Fructose bisphosphate aldolase protein was submitted to the following different B cell tests ^[54]^.

#### 2.3.1 Prediction of linear B-cell epitopes

Bepipred tool from IEDB (http://tools.iedb.org/bcell/result/) was used to predict the linear B-cell epitopes from the conserved region with a default threshold value of 0.350 ^[55–57]^.

#### 2.3.2 Prediction of surface accessibility

Emini surface accessibility prediction tool of the immune epitope database (IEDB) (http://tools.iedb.org/bcell/result/) was used to predict the surface epitopes from the conserved region with the default threshold value 1.0 ^[58]^

#### 2.3.3 Prediction of epitopes antigenicity

Kolaskar and Tongaonkar antigenicity method was used to detect the antigenic sites with a default threshold value of 1.025, (http://tools.iedb.org/bcell/result/ ^[59]^.

#### 2.3.4 Prediction of discontinuous B-cell Epitope

The modeled 3D structure was submitted to ElliPro prediction tool to filter out the antigenic residues. The minimum score and maximum distance (Angstrom) were calibrated in the default mode with a score of 0.5, and 6, respectively ^[60]^.

### 2.3 Prediction of MHC class I binding epitopes

The peptide binding affinity to MHC1 molecules was defined by the IEDB MHC I prediction tool at http://tools.iedb.org/mhc1. The binding affinity of Fructose bisphosphate aldolase peptide to MHC1 molecules was obtained using artificial neural network (ANN) method. All conserved epitopes that bind to MHC1 alleles at score ≤ 500 half-maximal inhibitory concentrations (IC50) with 9 peptides length were selected for further analysis ^[61–66]^.

### 2.5 MHC class II binding predictions

Prediction of peptide binding affinity to MHC11 molecules was defined by the IEDB MHC II prediction tool at (http://tools.iedb.org/mhcii/result/). MHC II molecule has the ability to bind different lengths Peptides which makes prediction accuracy debatable. For MHC II binding predication, human allele references set were used.

The prediction method was selected as NN-align to asses both the binding affinity and MHC II binding core epitopes with 9 peptides length at score IC50 of 100 ^[67]^.

### 2.6 Population coverage calculation

The candidates epitopes of MHC I and MHC II and combined binding of MHCI and MHC II alleles from *Candida glabrata* Fructose bisphosphate aldolase protein were employed for population coverage, and the world population was set as a target population for the selected MHC I and MHC II combined binding alleles using IEDB population coverage calculation tool at http://tools.iedb.org/population/ ^[68]^.

### 2.7 Homology modeling

The reference sequence of *Candida glabrata* Fructose bisphosphate aldolase protein was applied to Raptor X for modeling at: (http://raptorx.uchicago.edu/). Then the 3D structural model of the protein was visualized by using Chimera a tool powered by UCSF. ^[69–73]^.

### 2.8 The Physicochemical Properties

The function of vaccines is to enhance the immunogenic response once introduced to the immune system. Thus it is essential to recognize the physiochemical parameters of the protein protogram and Bio edit were implemented to assess the Physicochemical Properties. ^[53]^ (Available at: https://web.expasy.org/protparam/, and https://web.expasy.org/protscale/) ^[74]^.

### 2.9 Molecular docking analysis

Molecular docking was performed using Moe 2007. The 3D structures of the promiscuous epitopes were predicted by PEP-FOLD. The crystal structure of HLA A*02:06 (**PDB ID 3OXR**) and HLA-DRB1*01:01(**PDB ID 5JLZ**) were chosen as a model for molecular docking and were downloaded in a PDB format from the RCSB PDB resource. However, the selected crystal structures were in a complex form with ligands. Thus, to simplify the complex structure all water molecules, hetero groups and ligands were removed by Discovery Studio Visualizer 2.5. Partial charge and energy minimization were applied for ligands and targets. In terms of the identification of the binding groove, the potential binding sites in the crystal structure were recognized using the Alpha Site Finder. Finally, ten independent docking runs were carried out for each Peptide. The results were retrieved as binding energies. Best poses for each epitope that displayed lowest binding energies were visualized using UCSF chimera 1.13.1 software [72, 75–78]

## 3. Result

### 3.1.1 B-cell epitope prediction

The reference sequence of Fructose bisphosphate aldolase from *C. glabrata* was analyzed using Bepipred Linear Epitope Prediction first, the average binder’s score of the protein to B cell was 0.199, minimum was −0.009 and 2.424 for a maximum score, all values equal or greater than the default threshold 0.350 were potentially linear epitopes (Table 1 and figure 1).

**Table 1:** Predicted antigenic B-cell epitopes, 9 antigenic sites were identified from Fructose bisphosphate aldolase of *C. glabrata*.

| Start | End | Peptide | Length |
| --- | --- | --- | --- |
| **63** | 70 | SNGGAAYF | 8 |
| **73** | 84 | KGVSNDGQNASI | 12 |
| **129** | 134 | AYFKEH | 6 |
| **147** | 155 | SEETDDENI | 9 |
| **178** | 199 | ITGGEEDGVNNEHVDKESLYTK | 22 |
| **247** | 260 | KYAAEKTGAPAGSK | 14 |
| **269** | 280 | GSGSTQEEFNTG | 12 |
| **318** | 331 | GNPEGADKPNKKFF | 14 |
| **336** | 345 | WVREGEKTMS | 10 |

**Figure 1.**
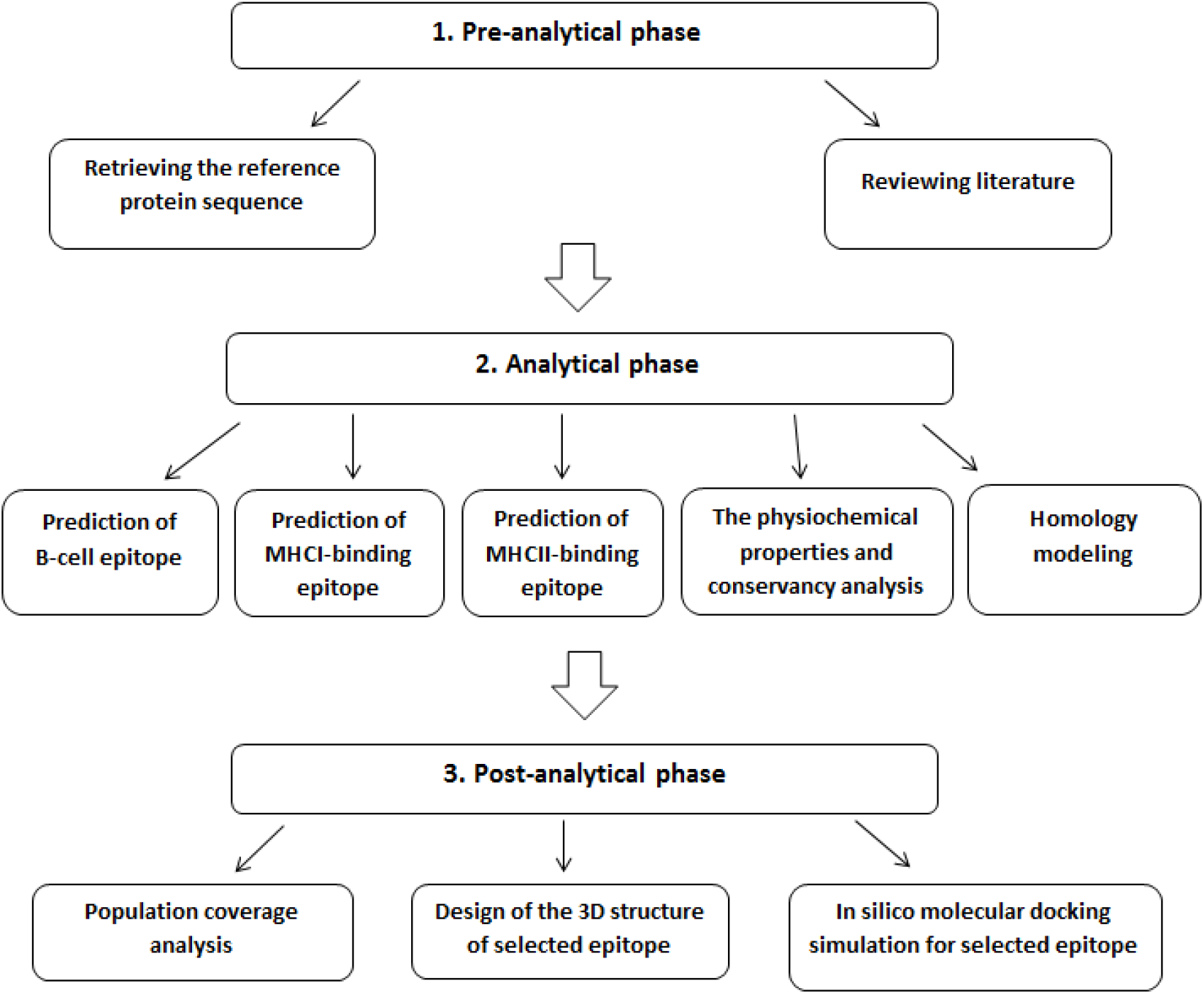
Schematic representation of the methodology phases

In Emini surface accessibility prediction, for a potent B-cell epitope the average surface accessibility area of the protein was scored as 1.000, with a maximum of 7.725 and a minimum of 0.113, all values equal or greater than the default threshold 1.000 were potentially in the surface. In addition, Kolaskar and Tongaonkar antigenicity prediction’s average of antigenicity was 1.025, with a maximum of 1.223 and minimum of 0.853; all values equal to or greater than the default threshold 1.025 are potential antigenic determinants. The results of all conserved predicted B cell epitopes are shown in (table 2 and figure 2).

**Table 2.**
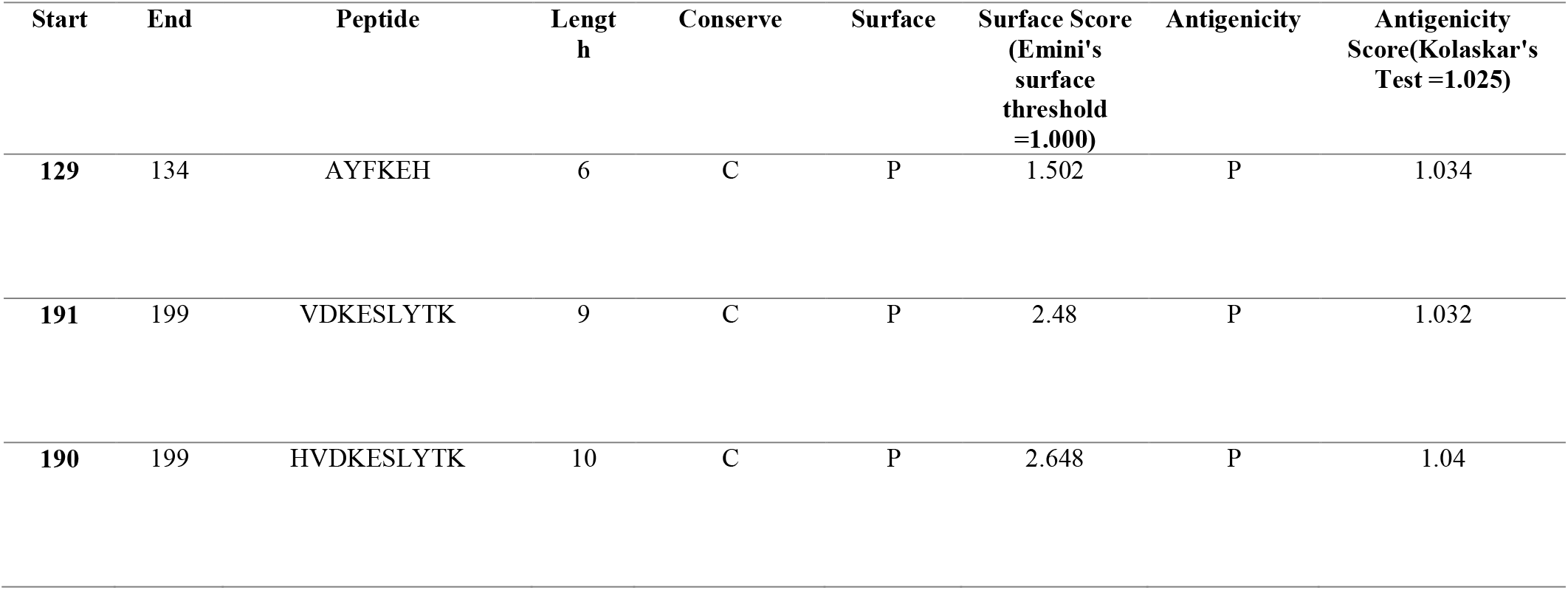
List of the most promising B-cell Epitope and their surface and Antigenicity (c=conserve, p=pathogenicity).

| Start | End | Peptide | Length | Conserve | Surface | Surface Score (Emini's surface threshold =1.000) | Antigenicity | Antigenicity Score(Kolaskar's Test =1.025) |
| --- | --- | --- | --- | --- | --- | --- | --- | --- |
| 129 | 134 | AYFKEH | 6 | C | P | 1.502 | P | 1.034 |
| 191 | 199 | VDKESLYTK | 9 | C | P | 2.48 | P | 1.032 |
| 190 | 199 | HVDKESLYTK | 10 | C | P | 2.648 | P | 1.04 |

**Figure 2:**
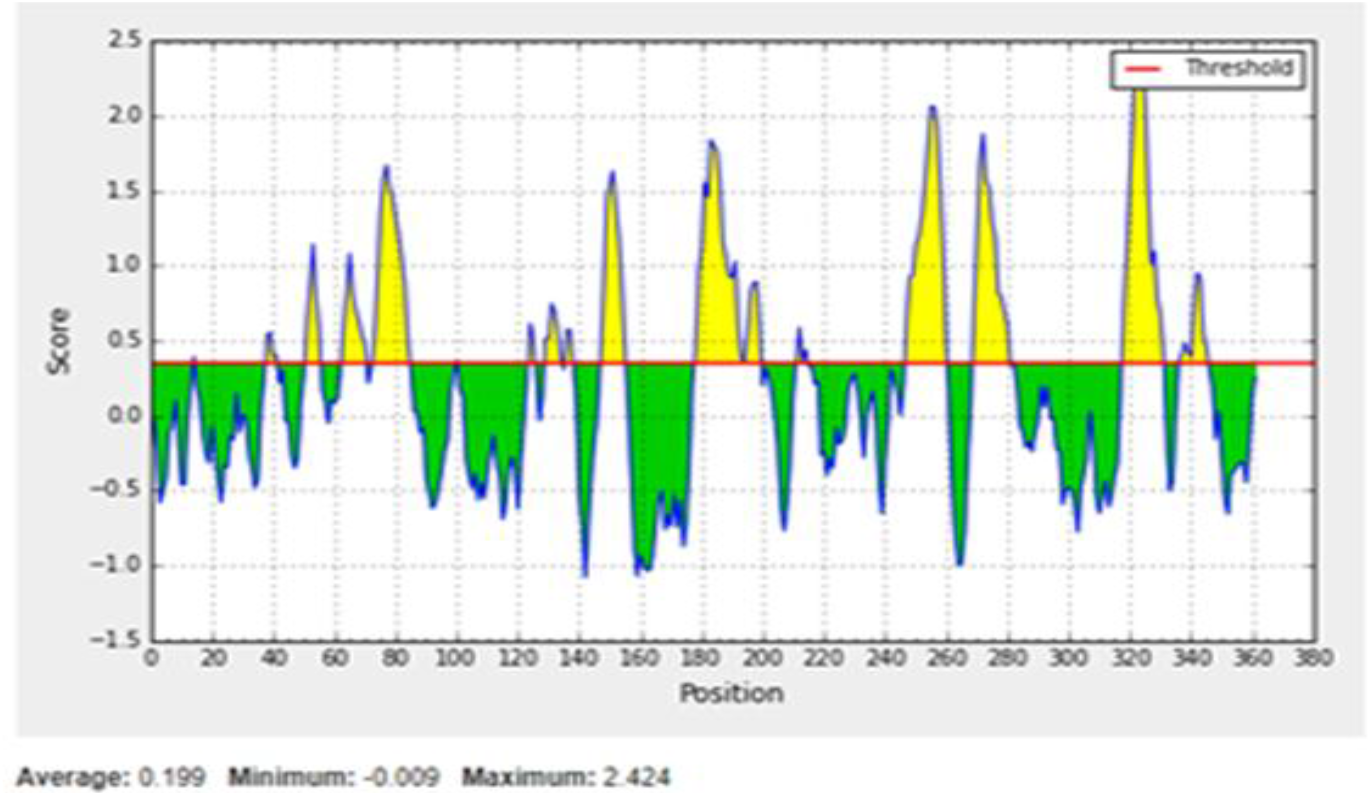
Bepipred Linear Epitope Prediction, The Red Line is the threshold, above (the yellow part) is proposed to be part of the B-cell Epitope.

**Figure 3:**
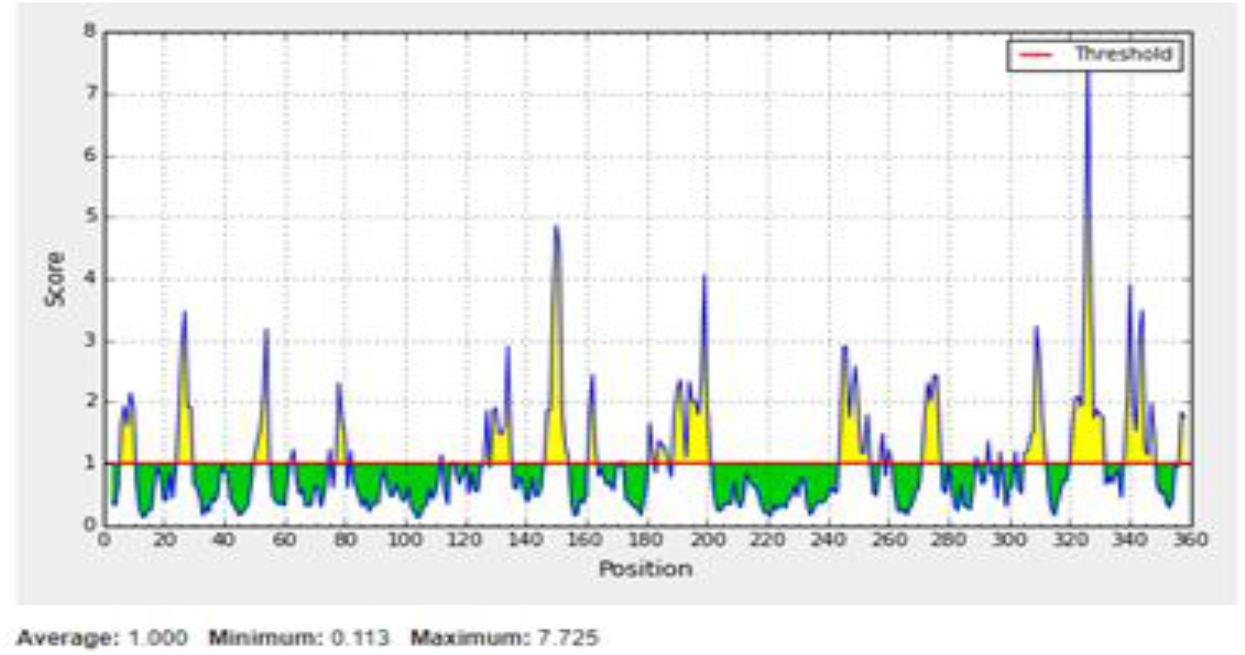
Emini’s Surface Accessibility Prediction Test, The Red Line is the threshold above (the yellow part) is proposed to be part of the B-cell Epitope.

### 3.1.2. Discontinuous B-cell Epitope Prediction

The modeled 3D structure was submitted to ElliPro prediction tool to filter out the antigenic residues. The minimum score and maximum distance (Angstrom) were calibrated in the default mode with a score of 0.5, and 6, respectively (Table3 and Figure 4).

**Table 3.**
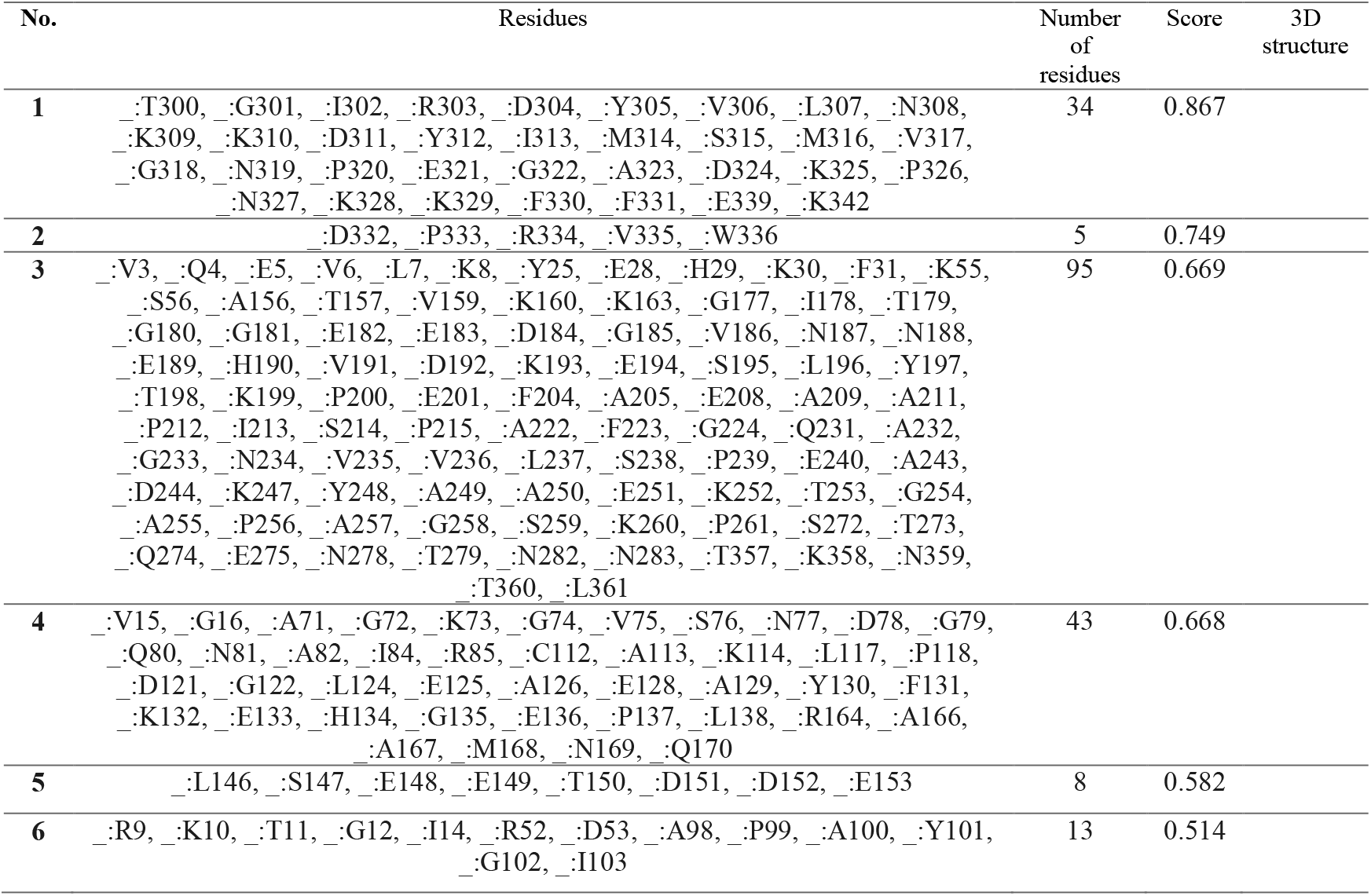
List of four predicted non-linear B-cell epitopes with highest number of Residues and their scores using ElliPro tool.

| No. | Residues | Number of residues | Score | 3D structure |
| --- | --- | --- | --- | --- |
| 1 | _:T300, _:G301, _:I302, _:R303, _:D304, _:Y305, _:V306, _:L307, _:N308, _:K309, _:K310, _:D311, _:Y312, _:I313, _:M314, _:S315, _:M316, _:V317, _:G318, _:N319, _:P320, _:E321, _:G322, _:A323, _:D324, _:K325, _:P326, _:N327, _:K328, _:K329, _:F330, _:F331, _:E339, _:K342 | 34 | 0.867 |  |

|  |  |  |  |
| --- | --- | --- | --- |
| 2 | _:D332, _:P333, _:R334, _:V335, _:W336 | 5 | 0.749 |
| 3 | _:V3, _:Q4, _:E5, _:V6, _:L7, _:K8, _:Y25, _:E28, _:H29, _:K30, _:F31, _:K55, _:S56, _:A156, _:T157, _:V159, _:K160, _:K163, _:G177, _:I178, _:T179, _:G180, _:G181, _:E182, _:E183, _:D184, _:G185, _:V186, _:N187, _:N188, _:E189, _:H190, _:V191, _:D192, _:K193, _:E194, _:S195, _:L196, _:Y197, _:T198, _:K199, _:P200, _:E201, _:F204, _:A205, _:E208, _:A209, _:A211, _:P212, _:I213, _:S214, _:P215, _:A222, _:F223, _:G224, _:Q231, _:A232, _:G233, _:N234, _:V235, _:V236, _:L237, _:S238, _:P239, _:E240, _:A243, _:D244, _:K247, _:Y248, _:A249, _:A250, _:E251, _:K252, _:T253, _:G254, _:A255, _:P256, _:A257, _:G258, _:S259, _:K260, _:P261, _:S272, _:T273, _:Q274, _:E275, _:N278, _:T279, _:N282, _:N283, _:T357, _:K358, _:N359, _:T360, _:L361 | 95 | 0.669 |
| 4 | _:V15, _:G16, _:A71, _:G72, _:K73, _:G74, _:V75, _:S76, _:N77, _:D78, _:G79, _:Q80, _:N81, _:A82, _:I84, _:R85, _:C112, _:A113, _:K114, _:L117, _:P118, _:D121, _:G122, _:L124, _:E125, _:A126, _:E128, _:A129, _:Y130, _:F131, _:K132, _:E133, _:H134, _:G135, _:E136, _:P137, _:L138, _:R164, _:A166, _:A167, _:M168, _:N169, _:Q170 | 43 | 0.668 |
| 5 | _:L146, _:S147, _:E148, _:E149, _:T150, _:D151, _:D152, _:E153 | 8 | 0.582 |
| 6 | _:R9, _:K10, _:T11, _:G12, _:I14, _:R52, _:D53, _:A98, _:P99, _:A100, _:Y101, _:G102, _:I103 | 13 | 0.514 |

**Figure 4:**
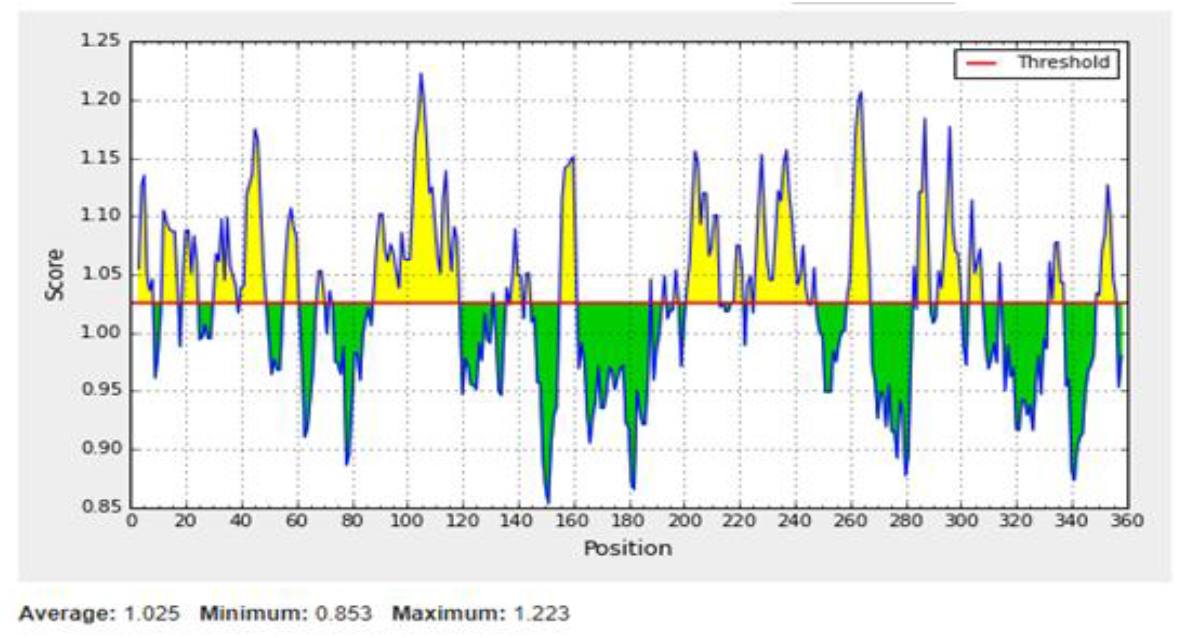
Kolaskar and Tongaonkar antigenicity prediction Test, the Red Line is the threshold, above (the yellow part) is proposed to be part of the B-cell Epitope.

### 3.2 T-cell peptide prediction

#### 3.2.1. Prediction of MHC-I binding profile for T cytotoxic cell conserved epitopes

114 epitopes were anticipated to interact with different MHC-1 alleles. The core epitopes (KYFKRMAAM and QTSNGGAAY) were noticed to be the dominant binders with 7 alleles for each (HLA-A*24:02, HLA-A*30:01, HLA-A*31:01, HLA-B*14:02, HLA-C*07:02, HLA-C*12:03, HLA-C*14:02) (HLA-A*01:01, HLA-A*26:01, HLA-A*29:02, HLA-A*30:02, HLA-B*15:01, HLA-B*15:02, HLA-B*35:01) followed by AVHEALAPI, RMAAMNQWL, and YFKEHGEPL which binds with five alleles (table IV).

**Table IV:** List of most promising epitopes that had a good binding affinity with MHC-I alleles in terms of IC50 and Percentile rank.

| Peptide | Start | End | Allele | ic50 |
| --- | --- | --- | --- | --- |
| **KYFKRMAAM** | 160 | 168 | HLA-A*24:02 | 451.84 |
|  | 160 | 168 | HLA-A*30:01 | 232.12 |
|  | 160 | 168 | HLA-A*31:01 | 131.22 |
|  | 160 | 168 | HLA-B*14:02 | 427.02 |
|  | 160 | 168 | HLA-C*07:02 | 149.13 |
|  | 160 | 168 | HLA-C*12:03 | 240.46 |
|  | 160 | 168 | HLA-C*14:02 | 6.27 |
| **AVHEALAPI** | 205 | 213 | HLA-A*02:01 | 154.37 |
|  | 205 | 213 | HLA-A*02:06 | 9.78 |
|  | 205 | 213 | HLA-A*30:01 | 20.96 |
|  | 205 | 213 | HLA-A*32:01 | 122.32 |
|  | 205 | 213 | HLA-A*68:02 | 55.22 |
| **RMAAMNQWL** | 164 | 172 | HLA-A*02:01 | 52.44 |
|  | 164 | 172 | HLA-A*02:06 | 237.09 |
|  | 164 | 172 | HLA-A*32:01 | 79.39 |
|  | 164 | 172 | HLA-B*15:01 | 258 |
|  | 164 | 172 | HLA-C*14:02 | 482 |
| **QTSNGGAAY** | 61 | 69 | HLA-A*01:01 | 54.18 |
|  | 61 | 69 | HLA-A*26:01 | 89.37 |
|  | 61 | 69 | HLA-A*29:02 | 56.68 |
|  | 61 | 69 | HLA-A*30:02 | 47.89 |
|  | 61 | 69 | HLA-B*15:01 | 111.57 |
|  | 61 | 69 | HLA-B*15:02 | 82.52 |
|  | 61 | 69 | HLA-B*35:01 | 99.45 |
| **YFKEHGEPL** | 130 | 138 | HLA-B*08:01 | 295.97 |
|  | 130 | 138 | HLA-C*03:03 | 42.03 |
|  | 130 | 138 | HLA-C*07:02 | 319.29 |
|  | 130 | 138 | HLA-C*12:03 | 26.8 |
|  | 130 | 138 | HLA-C*14:02 | 18.47 |

#### 3.2.2 Prediction of MHC-II binding profile for T helper cell conserved epitopes

102 conserved predicted epitopes were found to interact with MHC-II alleles. The core epitopes (LFSSHMLDL) is thought to be the top binder as it interacts with 9 alleles; (HLA-DRB1*07:01, HLA-DPA1*01, HLA-DPB1*04:01, HLA-DPA1*01:03, HLA-DPB1*02:01, HLA-DPA1*02:01, HLA-DPB1*01:01, HLA-DPA1*03:01, and HLA-DPB1*04:02) Followed by IRGSIAAAH which binds to five alleles and VVAALEAAR also bind with five alleles but low frequent. Followed by YQAGNVVLS, and IAPAYGIPV (Table V).

**Table V:**
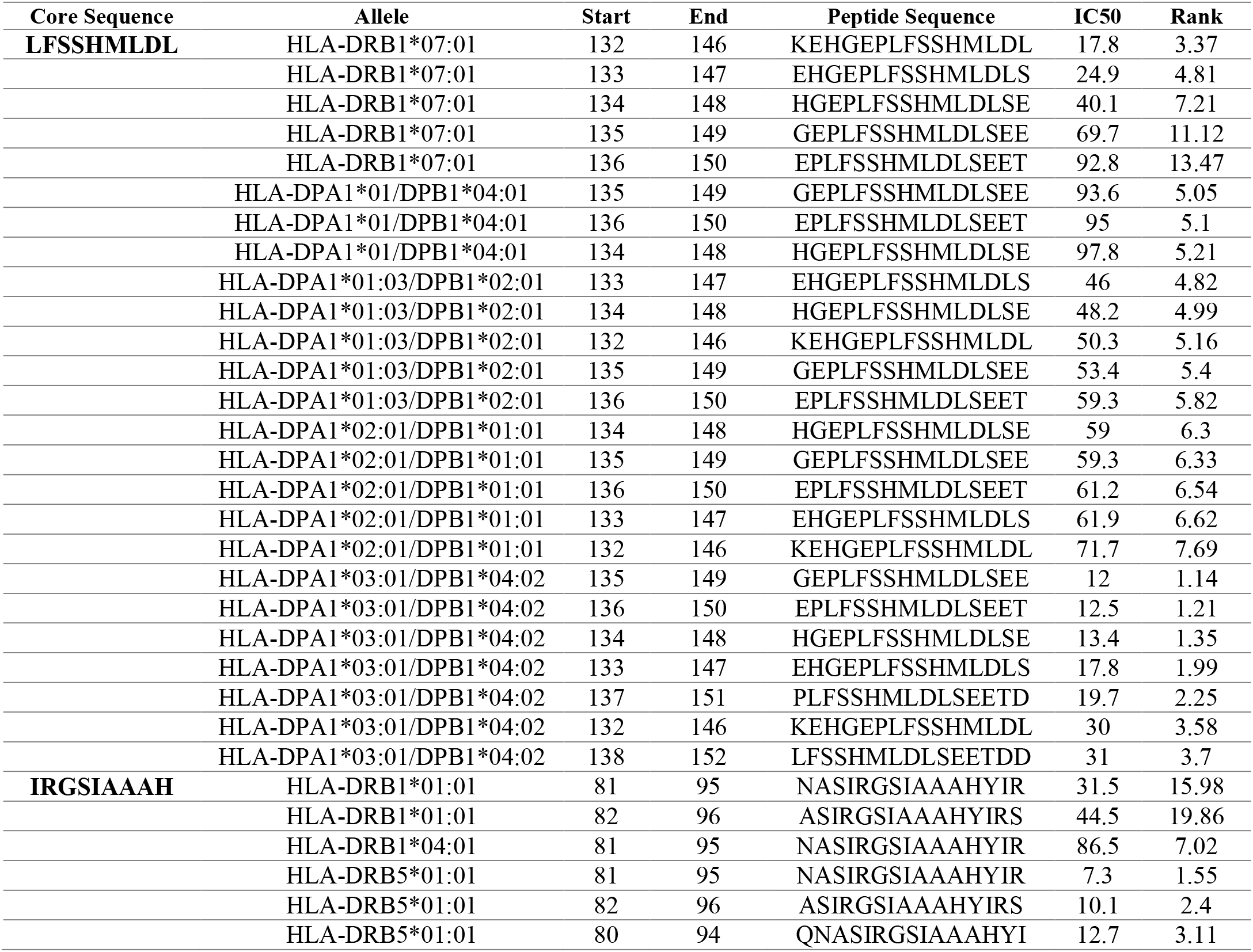
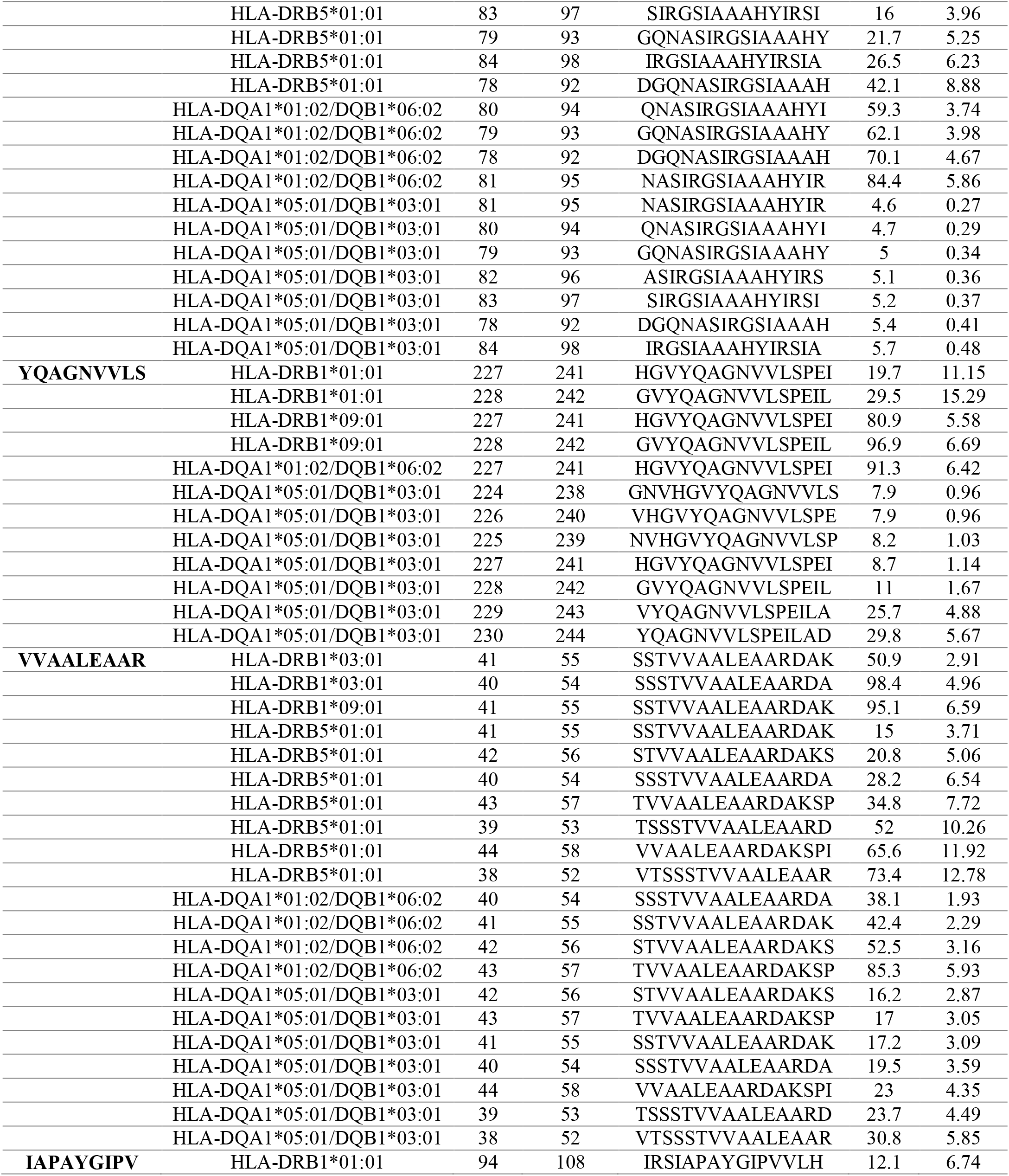
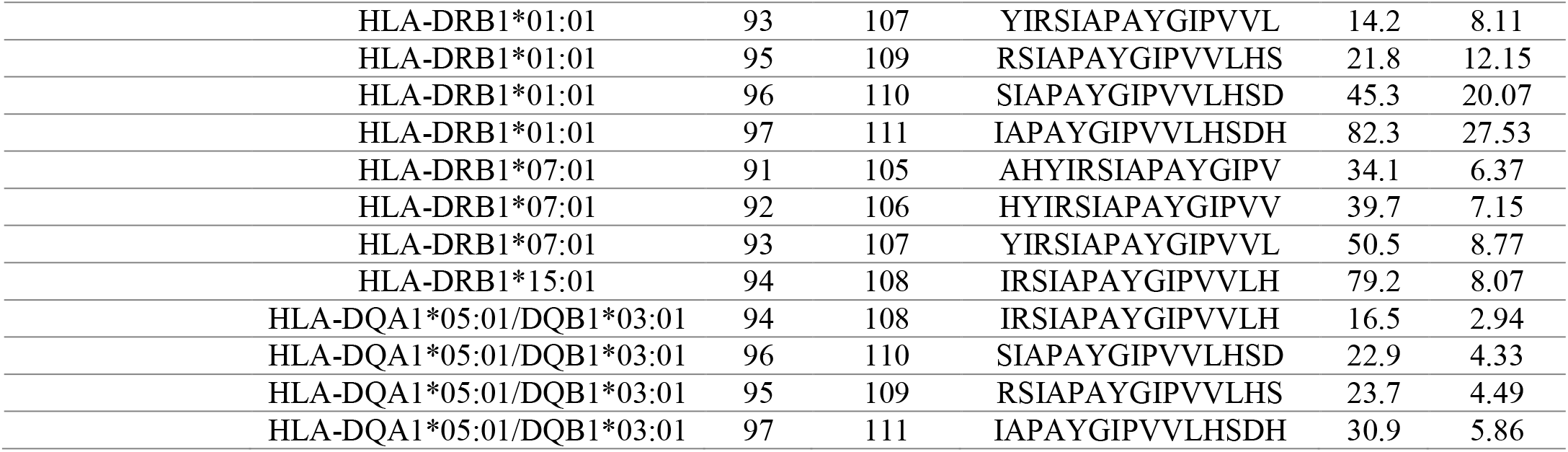
Promising T cell epitope (class MHC II Alleles) with their position and peptide sequence and ic50 value and rank

**Table VI:** Amino acids composition of the protein (Fructose-bisphosphate aldolase) with their number and molecular weight (Mol %) using Bio edit software version 7.0.5.3.

| Amino Acid | Number | Mol% | Amino Acid | Number | Mol% |
| --- | --- | --- | --- | --- | --- |
| Ala A | 39 | 10.80 | Leu L | 23 | 6.37 |
| Cys C | 3 | 0.83 | Met M | 9 | 2.49 |
| Asp D | 21 | 5.82 | Asn N | 19 | 5.26 |
| Glu E | 26 | 7.20 | Pro P | 15 | 4.16 |
| Phe F | 14 | 3.88 | Gln Q | 8 | 2.22 |
| Gly G | 30 | 8.31 | Arg R | 10 | 2.77 |
| His H | 12 | 3.32 | Ser S | 23 | 6.37 |
| Ile I | 20 | 5.54 | Thr T | 17 | 4.71 |
| Lys K | 25 | 6.93 | Val V | 29 | 8.03 |
| Trp W | 3 | 0.83 | Tyr Y | 15 | 4.16 |

### 3.3 Population coverage

The most interesting findings in this test is that the population coverage analysis result for most common binders to MHC-I and MHC-II alleles per each and combined; exhibit an exceptional coverage with percentages 92.54 %, 99.58 and 98,5 % Respectively.

#### 3.3.1 Population coverage for isolated MHC-I

Five epitopes are given to interact with the most frequent MHC class I alleles (**AVHEALAPI**, **KYFKRMAAM**, **QTSNGGAAY**, **RMAAMNQWL** and **YFKEHGEPL**), representing a considerable coverage against the whole world population. The maximum population coverage percentage over these epitopes is 92.54% (figure 5).

**Figure 5:**
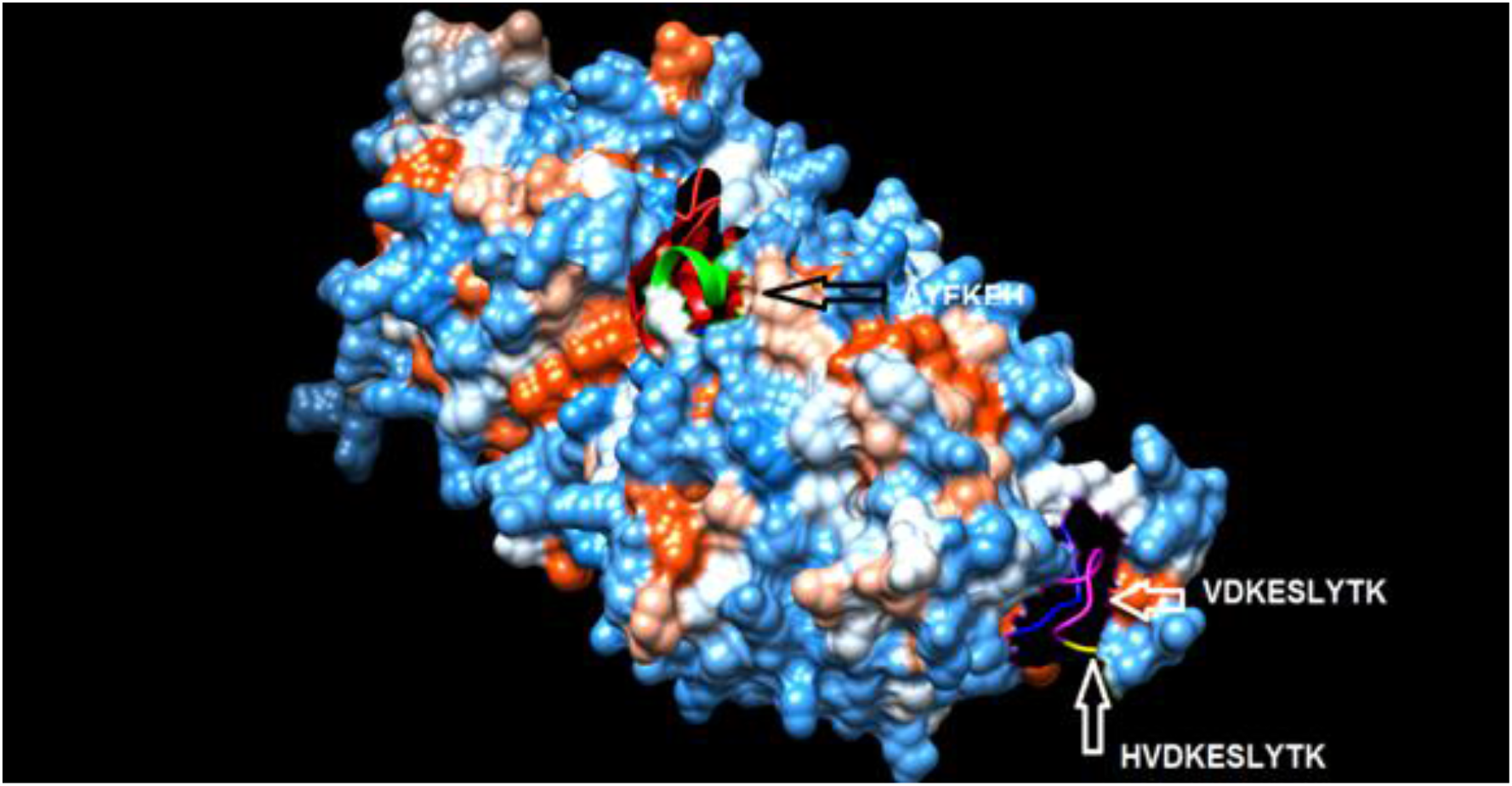
Structural position of the promising B-cell Epitope (AYFKPH, VDKESLYTK, and HVDKESLYTK).

#### 3.3.2 Population coverage for isolated MHC-II

Three epitopes were assumed to interact with the most frequent MHC class II alleles (IRGSIAAAH, LFSSHMLDL and VVAALEAAR) with percentage of 99.58%. The **LFSSHMLDL** epitope show an exceptional result for population coverage test for MHC II binding affinity of 96.60 % globally (figure 6).

**Figure 6.**
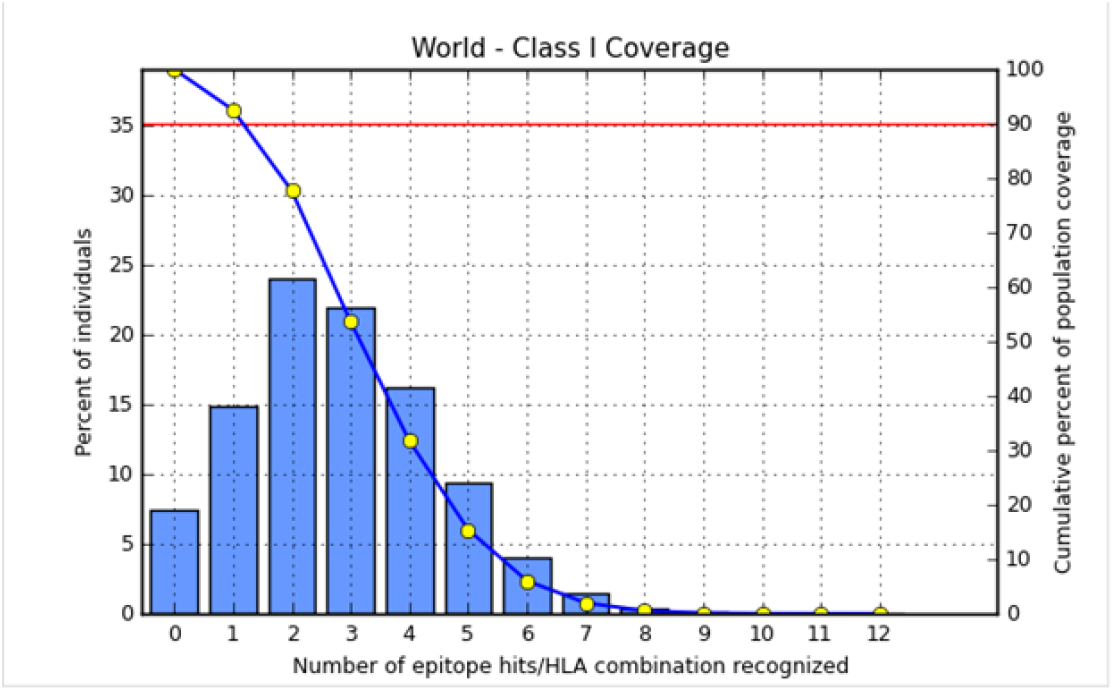
Illustrates the global coverage for the top five MHC-I peptides (AVHEALAPI, KYFKRMAAM, QTSNGGAAY, RMAAMNQWL and YFKEHGEPL) Note: In the graph, the line (-o-) represents the cumulative percentage of population coverage of the epitopes; the bars represent the population coverage for each epitope.

#### 3.3.3 Population coverage for MHC-I and MHC-II alleles combined

Regarding MHC-I and MHC-II alleles combined, five epitopes were supposed to interact with most predominant MHC class I and MHC class II alleles (IAPAYGIPV, AAFGNVHGV, VVAALEAAR, YIRSTIAPAY and YQAGMVVLS) represent a significant global coverage by IEDB population coverage tool. Reveal coverage with percentage of 98.50 % (figure 7).

**Figure 7.**
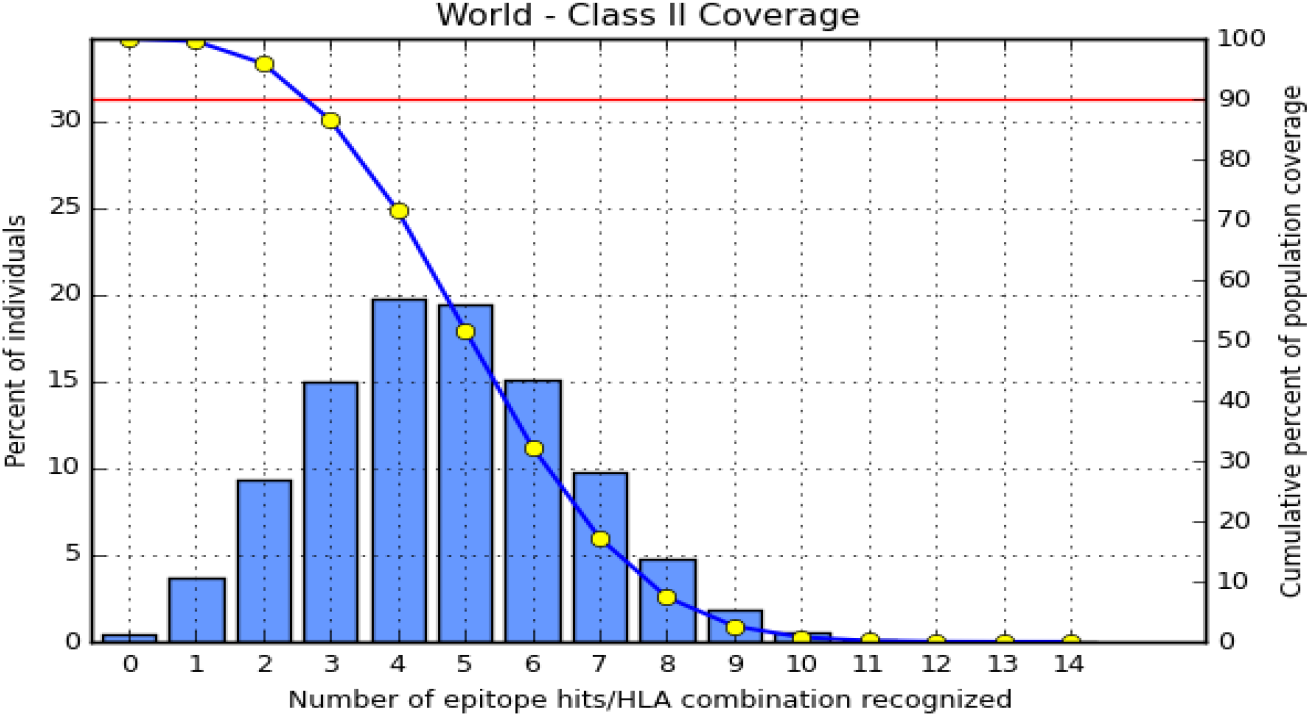
Illustrates the global proportion for the top five MHC-II IRGSIAAAH, LFSSHMLDL and VVAALEAAR. Notes: In the graph, the line (-o-) represents the cumulative percentage of population coverage of the epitopes; the bars represent the population coverage for each epitope.

### 3.4 Physiochemical properties

Predicted as 1 domain(s). Best template: 1dosA, p-value 4.00e-09. Overall uGDT (GDT): 299 (82). 361(100%) residues are modeled. 36(9%) positions predicted as disordered. Secondary struct: 44%H, 10%E, And 44% C. Solvent access: 34%E, 27%M, and 38% B.

### 3.5 Molecular docking

## 4. Discussion

In the present study, we predicted the most conserved and immunogenic B- and T-cell epitopes from Fba1 protein of *C. Glabrata* by using immunoinformatics approach in order to develop an effective epitope-based vaccine against this fungal pathogen which has emerged in recent years as serious health problem especially among immunosuppressed and hospitalized patient ^[79]^. In a previous study conducted by *de Klerk et al[80]*, showed that the Fba1 protein has the ability to provoke an immune responses in human against *M. mycetomatis*^[80]^. Also, several of recent publications have used the Fba1 protein as a strong antigenic target for predicting B- and T-cells epitopes in order to design promising vaccines against fungal and bacterial pathogens by using in silico tools such as *M. mycetomatis, P. Aeruginosa, L. Monocytogenes, and S. Mansoni* ^[81–84]^. Hence there are more studies to explore Fructose bisphosphate aldolase protein immunogenic role and the possibility to find common conserved epitopes for different organisms.

The principle of using a cocktail of B- and T-cell epitopes in the epitope-based vaccine to trigger humoral as well as cellular mediated immune responses is much promising to clear infection than humoral or cellular immunity alone, and it was applied before to enhance protection against different kind of infectious diseases ^[85, 86]^. In this study, the analysis of Fba1 protein revealed 11 effective epitopes for B cell (AYFKEH, VDKESLYTK and HVDKESLYTK) and T cell (AVHEALAPI, KYFKRMAAM, QTSNGGAAY, RMAAMNQWL, YFKEHGEPL, IRGSIAAAH, LFSSHMLDL and VVAALEAAR). However, the molecular docking, which evaluates the binding affinity to MHCs molecules ^[51, 52]^, showed that the peptides QTSNGGAAY and LFSSHMLDL are the best candidates for designing an effective epitope-based vaccine against *C. galabrata*.

After retrieving the various sequences of *C. glabrata* Fructose bisphosphate aldolase protein, the protein reference sequence was submitted to Bepipred linear epitope prediction test, Emini surface accessibility test and Kolaskar and Tongaonkar antigenicity test in the IEDB, to determine the affinity of B cell epitopes and their position regarding the surface and their immunogenicity. Three peptides have passed (AYFKEH, VDKESLYTK and HVDKESLYTK) in all the prediction tests (Table I, II) (Figure 2–4). However, the MHC I binding prediction tool using Artificial neural network (ANN) ^[61]^ with half-maximal inhibitory concentration (IC50) ≤ 500 revealed 114 conserved peptides interacting with various MHC I alleles. Three peptides were noticed to have the highest affinity in corresponding to their interaction with MHC I alleles. The peptide YIRSIAPAY from 93 to 101 had the affinity with 8 alleles to interact with (HLA-A*26:01,HLA-A*29:02,HLA-B *15:01,HLA-A*30:02,HLA-B *15:02,HLA-B*35:01,HLA-C*14:02,HLA-C*12:03), followed in order by KYFKRMAAM from 160 to 168 which interacts with 7 alleles (HLA-A*24:02,HLA-A*31:01,HLA-A*30:01,HLA-B*14:02,HLA-C*07:02,HLA-C*14:02,HLA-C*12:03) and QTSNGGAAY from 61 to 69 interacts with 7 alleles (HLA-A*01:01,HLA-A*26:01,HLA-A*30:02,HLA-A*29:02,HLA-B*15:02,HLA-B*15:01,HLA-B*35:01) (Table IV). While MHC II binding prediction tool using (NN-align) ^[67]^ with half-maximal inhibitory concentration (IC50) ≤ 100 revealed 102 conserved peptides that interact with various MHC II alleles. Two peptides (LFSSHMLDL and YIRSIAPAY) were noted to have the highest affinity in corresponding to their interaction with MHC II alleles, both had the affinity to interact with 9 MHC II alleles (Table V).

Moreover, the predicted epitopes which have the high affinity to interact with MHC I, MHC II and combined MHC I with MHC II international alleles were analyzed by population coverage resource in the IEDB ^[68]^. The population coverage of the five most promising epitopes (AVHEALAPI, KYFKRMAAM, QTSNGGAAY, RMAAMNQWL and YFKEHGEPL) for MHC I alleles was 92.54%, while for the three epitopes (IRGSIAAAH, LFSSHMLDL and VVAALEAAR) that showed high affinity to MHCII alleles was 99.58% throughout the world according to IEDB database (Figures 6–7). It should be noted that the population coverage of the five most promising epitopes that exhibited binding affinity to both MHC I and MHC II alleles (IAPAYGIPV, AAFGNVHGV, VVAALEAAR, YIRSTIAPAY and YQAGMVVLS) was 98.50% globally (Figure 8). However, the molecular docking revealed the epitopes QTSNGGAAY and LFSSHMLDL have high binding energy to MHC molecules HLA A*02:06 and HLA-DRB1*01:01, respectively which indicate favored affinity and stability in the epitope-molecule complex (Table VII) (Figure 10–17). This study was limited by being strictly computational; and more in vitro and in vivo studies to prove the effectiveness of the proposed peptides are highly recommended.

**Figure 8:**
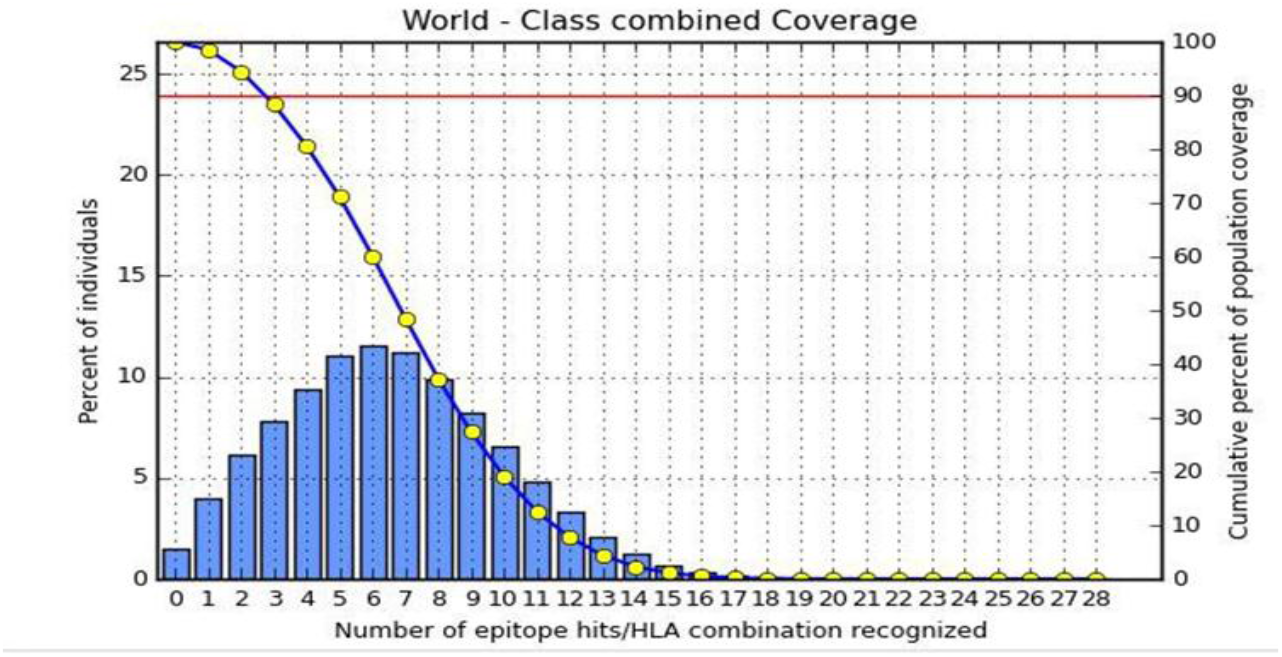
Illustrates the global population proportion for the top five MHC-I & II epitopes in combined mode (IAPAYGIPV, AAFGNVHGV, VVAALEAAR, YIRSTIAPAY and YQAGMVVLS). Notes: In the graphs, the line (-o-) represents the cumulative percentage of population coverage of the epitopes; the bars represent the population coverage for each epitope.

**Figure 9:**
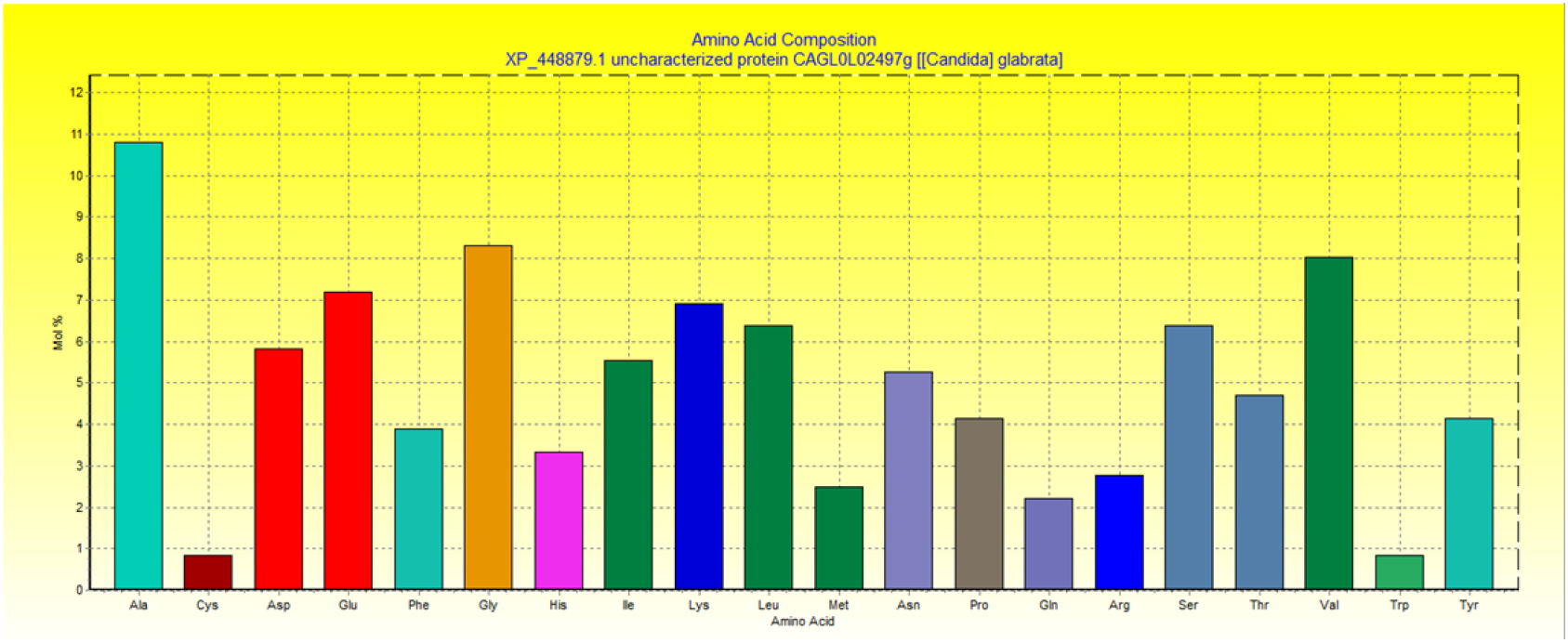
Graph showing amino acid composition of Fructose bisphosphate aldolase protein and their molecular weight using Bio edit software 7.0.5.3

**Table VII.** Docking results of the most promiscuous epitopes that show the best binding affinity

| Epitope | Binding MHC molecule | Binding Energy (*$\Delta G$ kcal/mol) |
| --- | --- | --- |
| AVHEALAPI | HLA A*02:06 | -15.8010 |
| KYFKRMAAM | HLA A*02:06 | -20.5935 |
| QTSNGGAAY | HLA A*02:06 | -30.5467 |
| RMAAMNQWL | HLA A*02:06 | -20.6392 |
| YFKEHGEPL | HLA A*02:06 | -16.7505 |
| IRGSIAAAH | HLA-DRB1*01:01 | -20.6557 |
| LFSSHMLDL | HLA-DRB1*01:01 | -25.5732 |
| VVAALEAAR | HLA-DRB1*01:01 | -19.8404 |

\*Global energy, it is the energy required to estimate the strength of association between the epitope within the active cleft of MHC molecule; more negative value indicates favored affinity and stable binding of the complex.

**Figure 10.**
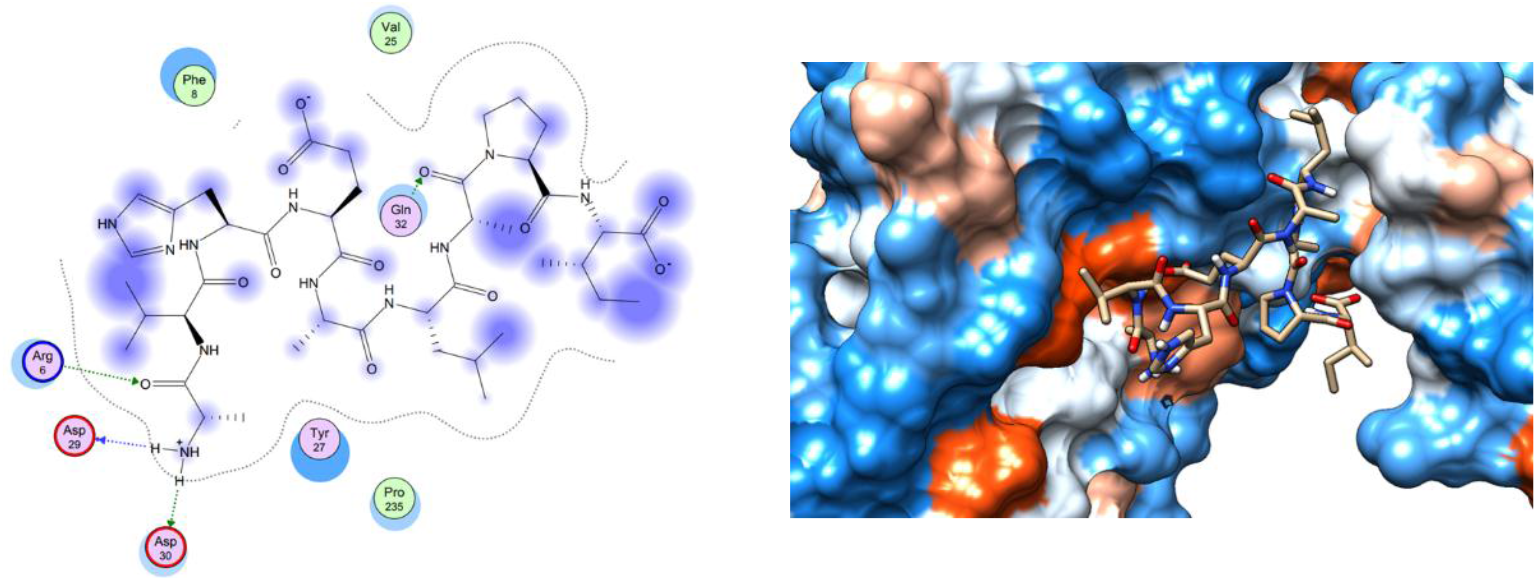
Illustrate the 2D interaction of the best docking poses of AVHEALAPI in the binding sites of HLA A*02:06.

**Figure 11.**
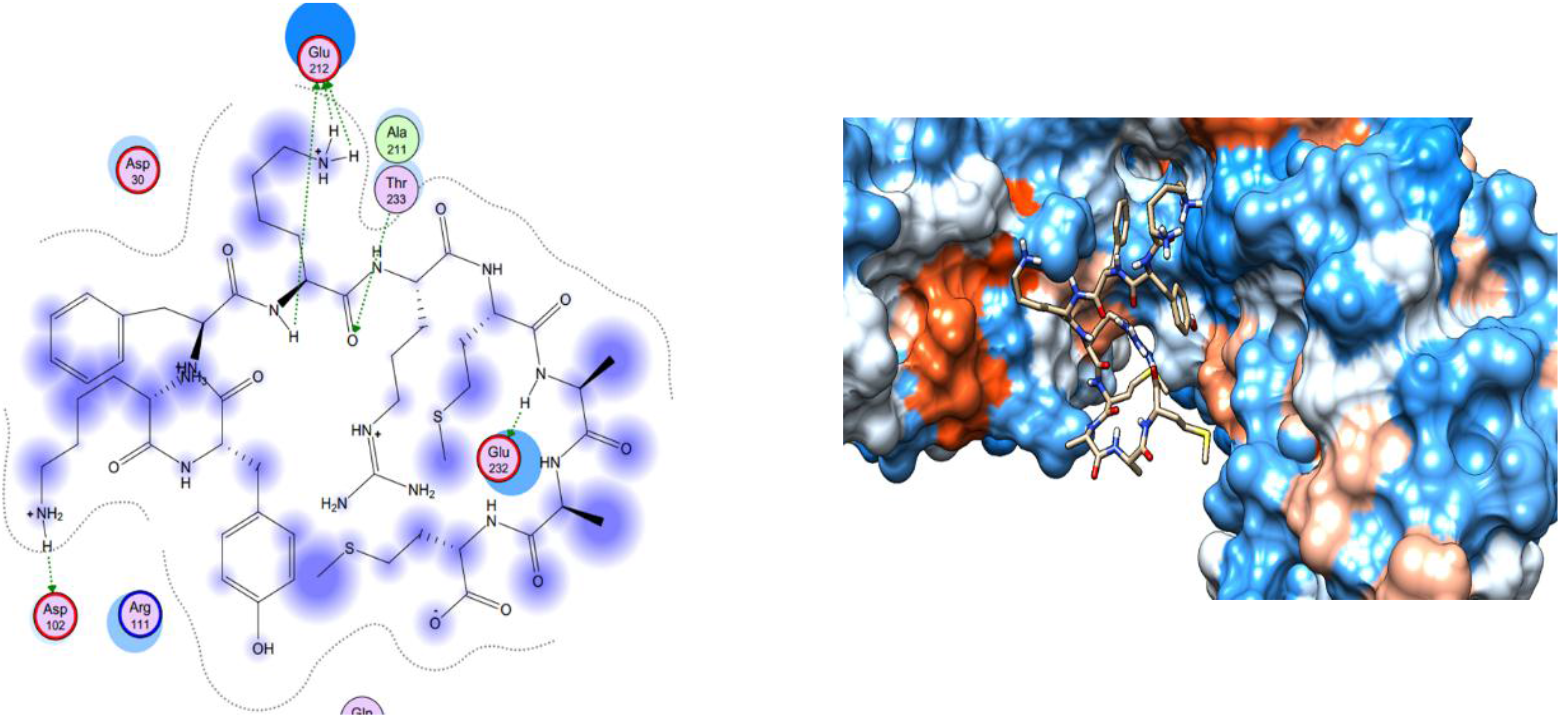
Illustrate the 3D interaction of the best docking poses of KYFKRMAAM in the binding sites of HLA A*02:06.

**Figure 12.**
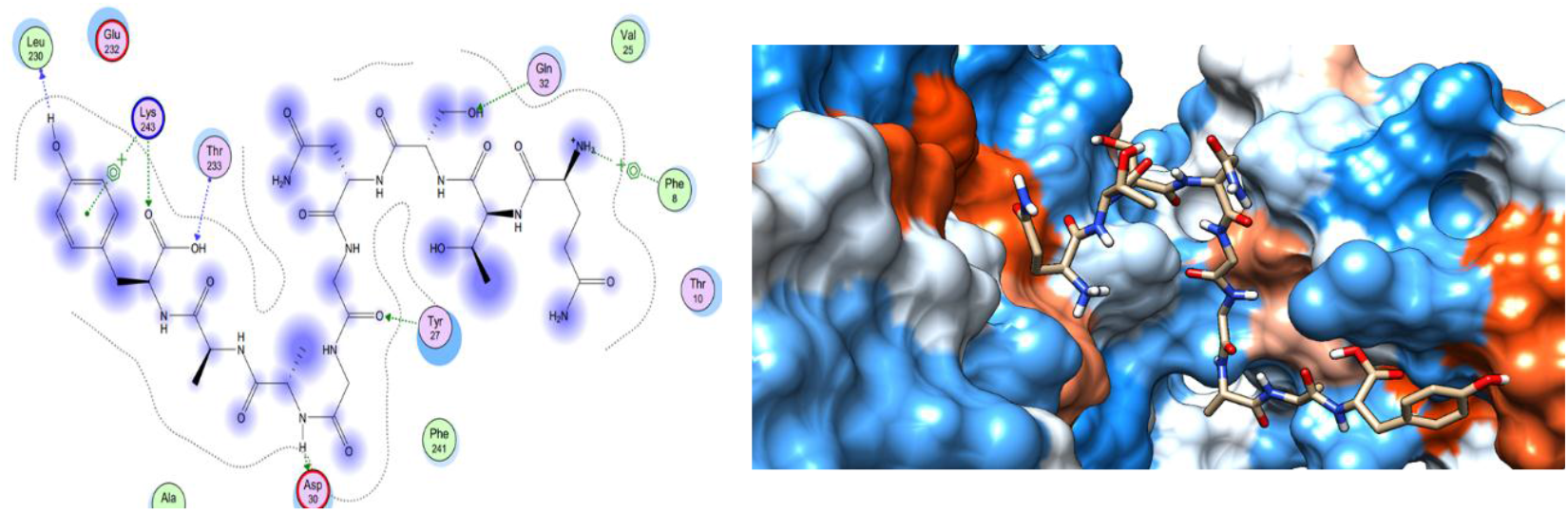
Illustrate the 2D interaction of the best docking poses of QTSNGGAAY in the binding sites of HLA A*02:06.

**Figure 13.**
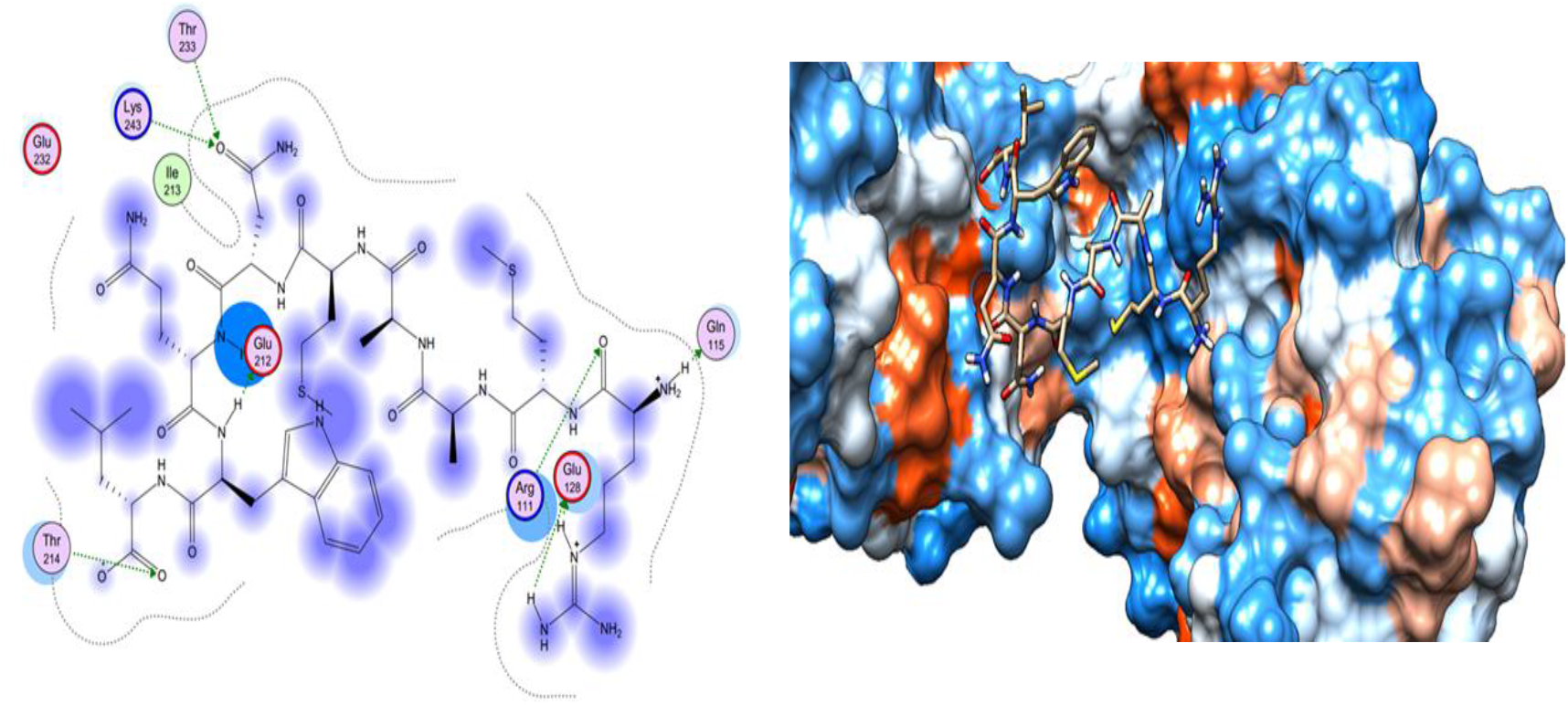
Illustrate the 2D interaction of the best docking poses of RMAAMNQWL in the binding sites of HLA A*02:06.

**Figure 14.**
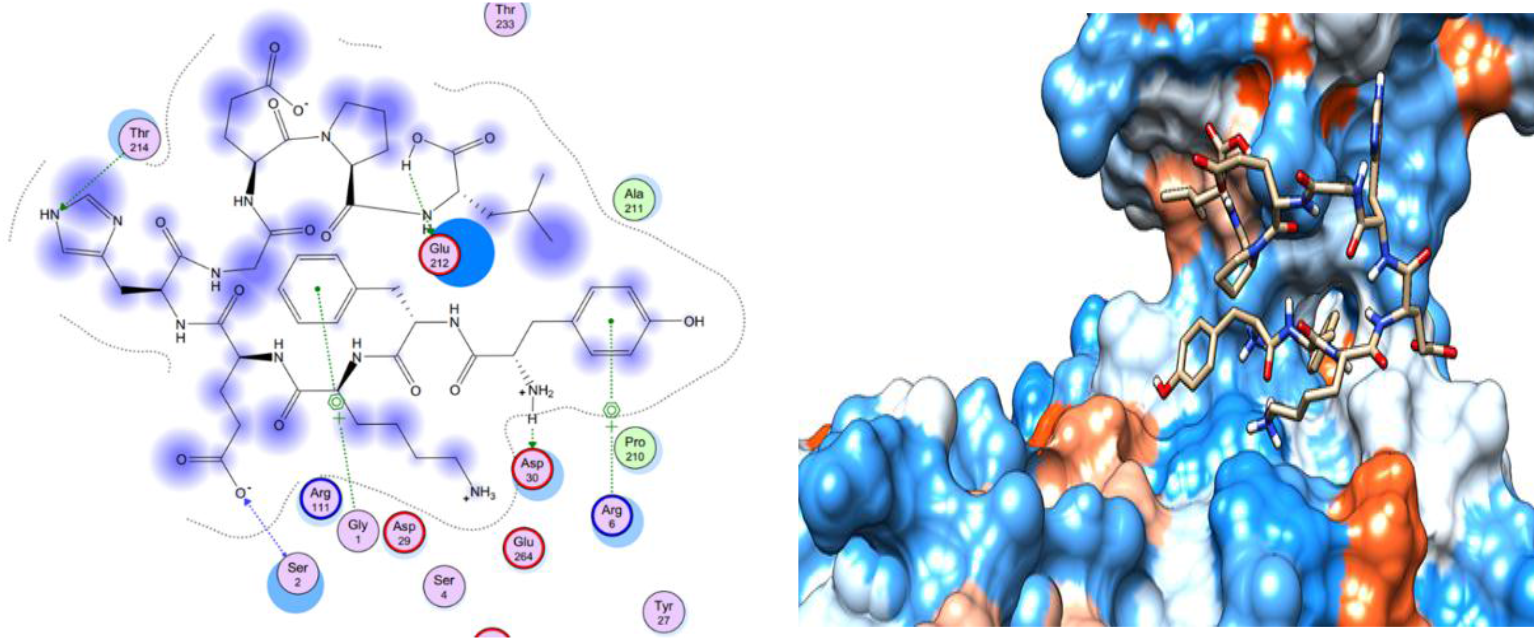
Illustrate the 3D interaction of the best docking poses of YFKEHGEPL in the binding sites of HLA A*02:06.

**Figure 15.**
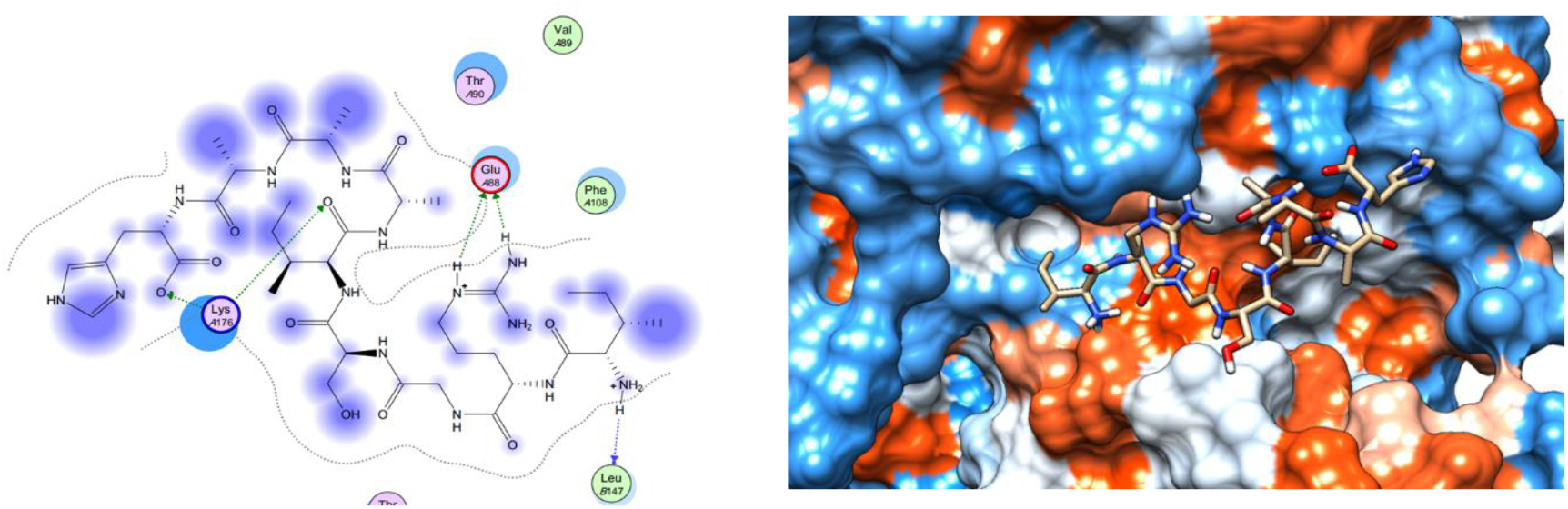
Illustrate the 3D interaction of the best docking poses of IRGSIAAAH in the binding sites of HLA-DRB1*01:01.

**Figure 16.**
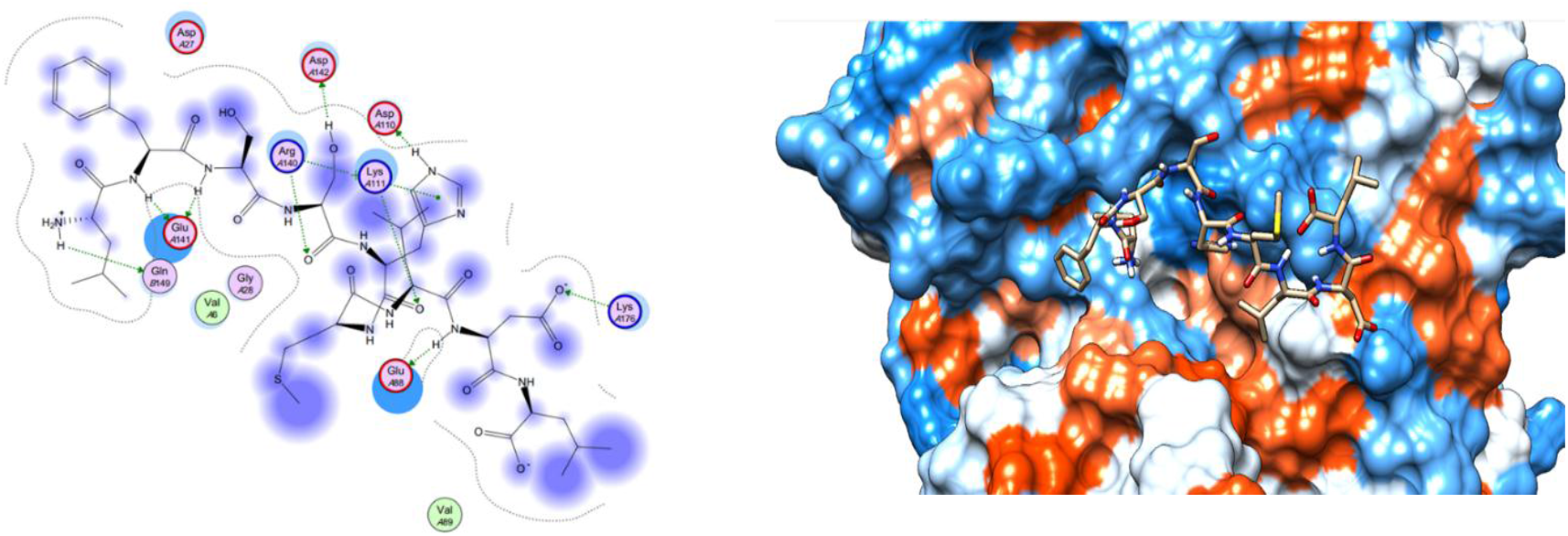
Illustrate the 3D interaction of the best docking poses of LFSSHMLDL the binding sites of HLA-DRB1*01:01.

**Figure 17.**
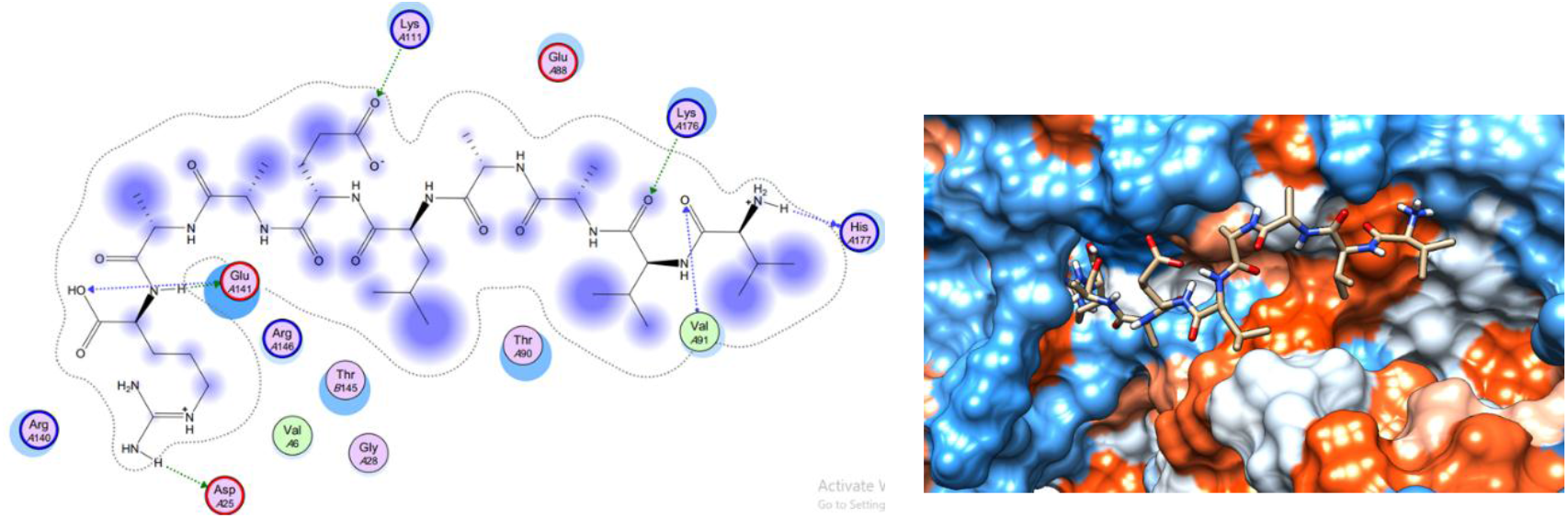
Illustrate the 2D interaction of the best docking poses of VVAALEAAR in the binding sites of HLA-DRB1*01:01.

## In conclusion

The epitope-based vaccines predicted by using immunoinformatics tools have remarkable advantages over the conventional vaccines that they are more specific, less time consuming, safe, less allergic and more antigenic. Further in vivo and in vitro experiments are needed to prove the effectiveness of the best candidate’s epitopes (QTSNGGAAY and LFSSHMLDL). To the best of our knowledge, this is the first study that has predicted B- and T-cells epitopes from Fba1 protein by using in silico tools in order to design an effective epitope-based vaccine against *C. galabrata*.

## Acknowledgment

The authors are grateful to Africa City of Technology, Sudan, Khartoum.

## Supplementary material

All data are included in this published article.

## References

1. Bitar, D., et al., Population-based analysis of invasive fungal infections, France, 2001-2010. Emerg Infect Dis, 2014. 20(7): p. 1149–55.

2. Cleveland, A.A., et al., Changes in incidence and antifungal drug resistance in candidemia: results from population-based laboratory surveillance in Atlanta and Baltimore, 2008-2011. Clin Infect Dis, 2012. 55(10): p. 1352–61.

3. Kullberg, B.J. and M.C. Arendrup, Invasive Candidiasis. N Engl J Med, 2015. 373(15): p. 1445–56.

4. Magill, S.S., et al., Multistate point-prevalence survey of health care-associated infections. N Engl J Med, 2014. 370(13): p. 1198–208.

5. Silva, S., et al., Candida glabrata, Candida parapsilosis and Candida tropicalis: biology, epidemiology, pathogenicity and antifungal resistance. FEMS Microbiology Reviews, 2012. 36(2): p. 288–305.

6. Wormser, G.P. and K.J. Ryan, Medically Important Fungi: A Guide to Identification, 4th Edition Davise H. Larone Washington, D.C.: American Society for Microbiology Press, 2002. 409 pp., illustrated. $79.95 (cloth). Clinical Infectious Diseases, 2003. 37(9): p. 1281–1281.

7. Brandt, M.E., Candida and Candidiasis. Emerging Infectious Diseases, 2002. 8(8): p. 876–876.

8. Kasper, L., K. Seider, and B. Hube, Intracellular survival of Candida glabrata in macrophages: immune evasion and persistence. FEMS Yeast Res, 2015. 15(5): p. fov042.

9. Atanasova, R., et al., A mouse model for Candida glabrata hematogenous disseminated infection starting from the gut: evaluation of strains with different adhesion properties. PLoS One, 2013. 8(7): p. e69664.

10. Dujon, B., et al., Genome evolution in yeasts. Nature, 2004. 430(6995): p. 35–44.

11. López-Fuentes, E., et al., Candida glabrata’s Genome Plasticity Confers a Unique Pattern of Expressed Cell Wall Proteins. Journal of fungi (Basel, Switzerland), 2018. 4(2): p. 67.

12. Ahmad, K.M., et al., Genome structure and dynamics of the yeast pathogen Candida glabrata. FEMS yeast research, 2014. 14(4): p. 529–535.

13. Gabaldón, T., et al., Comparative genomics of emerging pathogens in the Candida glabrata clade. BMC Genomics, 2013. 14(1): p. 623.

14. de Groot, P.W.J., et al., The cell wall of the human pathogen Candida glabrata: differential incorporation of novel adhesin-like wall proteins. Eukaryotic cell, 2008. 7(11): p. 1951–1964.

15. Timmermans, B., et al., Adhesins in Candida glabrata. Journal of fungi (Basel, Switzerland), 2018. 4(2): p. 60.

16. Nunez-Beltran, A., E. Lopez-Romero, and M. Cuellar-Cruz, Identification of proteins involved in the adhesionof Candida species to different medical devices. Microb Pathog, 2017. 107: p. 293–303.

17. Sun, X., et al., Association between IL-1b polymorphisms and gastritis risk. Medicines, 2017. 96: p. 5.

18. Martinez-Gomariz, M., et al., Proteomic analysis of cytoplasmic and surface proteins from yeast cells, hyphae, and biofilms of Candida albicans. Proteomics, 2009. 9(8): p. 2230–52.

19. Cabezon, V., et al., Apoptosis of Candida albicans during the Interaction with Murine Macrophages: Proteomics and Cell-Death Marker Monitoring. J Proteome Res, 2016. 15(5): p. 1418–34.

20. Leach, M.D., et al., Molecular and proteomic analyses highlight the importance of ubiquitination for the stress resistance, metabolic adaptation, morphogenetic regulation and virulence of Candida albicans. Mol Microbiol, 2011. 79(6): p. 1574–93.

21. Lee, K.H., et al., Proteomic analysis of hyphae-specific proteins that are expressed differentially in cakem1/cakem1 mutant strains of Candida albicans. J Microbiol, 2010. 48(3): p. 365–71.

22. Medrano-Diaz, C.L., et al., Moonlighting proteins induce protection in a mouse model against Candida species. Microb Pathog, 2018. 124: p. 21–29.

23. Fell, D.A., Metabolic control analysis: a survey of its theoretical and experimental development. The Biochemical journal, 1992. 286 (Pt 2)(Pt 2): p. 313–330.

24. Schwelberger, H.G., S.D. Kohlwein, and F. Paltauf, Molecular cloning, primary structure and disruption of the structural gene of aldolase from Saccharomyces cerevisiae. Eur J Biochem, 1989. 180(2): p. 301–8.

25. Marsh, J. and H. Lebherz, Fructose-bisphosphate aldolases: An evolutionary history. Trends in biochemical sciences, 1992. 17: p. 110–3.

26. Compagno, C., B. Ranzi, and E. Martegani, The promoter of Saccharomyces cerevisiae FBA1 gene contains a single positive upstream regulatory element. FEBS letters, 1991. 293: p. 97–100.

27. Gao, X., et al., Fructose-1,6-bisphosphate aldolase of Mycoplasma bovis is a plasminogen-binding adhesin. Microb Pathog, 2018. 124: p. 230–237.

28. Ramírez-Quijas, M.D., E. López-Romero, and M. Cuéllar-Cruz, Proteomic analysis of cell wall in four pathogenic species of Candida exposed to oxidative stress. Microbial Pathogenesis, 2015. 87: p. 1–12.

29. Serrano Fujarte, I. and M. Cuéllar Cruz, Moonlight-like proteins of the cell wall protect sessile cells of Candida from oxidative stress. Microbial pathogenesis, 2016. 90.

30. Klis, F., et al., Covalently linked cell wall proteins of Candida albicans and their role in fitness and virulence. FEMS yeast research, 2009. 9: p. 1013–28.

31. Núñez-Beltrán, A. and M. Cuéllar Cruz, Identification of proteins involved in the adhesionof Candida species to different medical devices. Microbial Pathogenesis, 2017. 107: p. 293–303.

32. Laín, A., et al., Use of Recombinant Antigens for the Diagnosis of Invasive Candidiasis. Clinical and Developmental Immunology, 2008. 2008: p. 721950.

33. Jong, A.Y., et al., Binding of Candida albicans enolase to plasmin(ogen) results in enhanced invasion of human brain microvascular endothelial cells. Journal of Medical Microbiology, 2003. 52(8): p. 615–622.

34. Sandini, S., et al., A highly immunogenic recombinant and truncated protein of the secreted aspartic proteases family (rSap2t) of Candida albicans as a mucosal anticandidal vaccine. FEMS Immunology & Medical Microbiology, 2011. 62(2): p. 215–224.

35. Serrano-Fujarte, I., E. López-Romero, and M. Cuéllar-Cruz, Moonlight-like proteins of the cell wall protect sessile cells of Candida from oxidative stress. Microbial pathogenesis, 2015. 90: p. 22–33.

36. Rodaki, A., T. Young, and A.J.P. Brown, Effects of depleting the essential central metabolic enzyme fructose-1,6-bisphosphate aldolase on the growth and viability of Candida albicans: implications for antifungal drug target discovery. Eukaryotic cell, 2006. 5(8): p. 1371–1377.

37. Elhaik Goldman, S., et al., Streptococcus pneumoniae fructose-1,6-bisphosphate aldolase, a protein vaccine candidate, elicits Th1/Th2/Th17-type cytokine responses in mice. Int J Mol Med, 2016. 37(4): p. 1127–38.

38. Huang, J., et al., Fructose-1,6-bisphosphate aldolase is involved in Mycoplasma bovis colonization as a fibronectin-binding adhesin. Res Vet Sci, 2019. 124: p. 70–78.

39. Wilde, S., et al., Salmonella-vectored vaccine delivering three Clostridium perfringens antigens protects poultry against necrotic enteritis. PLoS One, 2019. 14(2): p. e0197721.

40. Ahmad, K.M., et al., Genome structure and dynamics of the yeast pathogen Candida glabrata. FEMS Yeast Res, 2014. 14(4): p. 529–35.

41. Brunke, S. and B. Hube, Two unlike cousins: Candida albicans and C. glabrata infection strategies. Cell Microbiol, 2013. 15(5): p. 701–8.

42. Healey, K.R. and D.S. Perlin, Fungal Resistance to Echinocandins and the MDR Phenomenon in Candida glabrata. J Fungi (Basel), 2018. 4(3).

43. Wilson, L.S., et al., The direct cost and incidence of systemic fungal infections. Value Health, 2002. 5(1): p. 26–34.

44. Clerckx, C., et al., Candida glabrata renal abscesses in a peritoneal dialysis patient. Perit Dial Int, 2012. 32(1): p. 114–5.

45. Eschenauer, G.A., et al., Survival in Patients with Candida glabrata Bloodstream Infection Is Associated with Fluconazole Dose. Antimicrob Agents Chemother, 2018. 62(6).

46. Timmermans, B., et al., Adhesins in Candida glabrata. J Fungi (Basel), 2018. 4(2).

47. Zhu, Z., et al., Multiple brain abscesses caused by infection with Candida glabrata: A case report. Exp Ther Med, 2018. 15(3): p. 2374–2380.

48. Nami, S., et al., Fungal vaccines, mechanism of actions and immunology: A comprehensive review. Biomed Pharmacother, 2019. 109: p. 333–344.

49. Xin, H., Effects of immune suppression in murine models of disseminated Candida glabrata and Candida tropicalis infection and utility of a synthetic peptide vaccine. Med Mycol, 2018.

50. He, Y., et al., Vaccine informatics. J Biomed Biotechnol, 2010. 2010: p. 765762.

51. Backert, L. and O. Kohlbacher, Immunoinformatics and epitope prediction in the age of genomic medicine. Genome Med, 2015. 7: p. 119.

52. Bahrami, A.A., et al., Immunoinformatics: In Silico Approaches and Computational Design of a Multi-epitope, Immunogenic Protein. Int Rev Immunol, 2019. 38(6): p. 307–322.

53. Hall, T.A. BioEdit: a user-friendly biological sequence alignment editor and analysis program for Windows 95/98/NT. in Nucleic acids symposium series. 1999. [London]: Information Retrieval Ltd., c1979–c2000.

54. Vita, R., et al., The immune epitope database (IEDB) 3.0. Nucleic Acids Res, 2015. 43(Database issue): p. D405–12.

55. Haste Andersen, P., M. Nielsen, and O. Lund, Prediction of residues in discontinuous B-cell epitopes using protein 3D structures. Protein Sci, 2006. 15(11): p. 2558–67.

56. Larsen, J.E., O. Lund, and M. Nielsen, Improved method for predicting linear B-cell epitopes. Immunome Res, 2006. 2: p. 2.

57. Ponomarenko, J.V. and P.E. Bourne, Antibody-protein interactions: benchmark datasets and prediction tools evaluation. BMC Struct Biol, 2007. 7: p. 64.

58. Emini, E.A., et al., Induction of hepatitis A virus-neutralizing antibody by a virus-specific synthetic peptide. J Virol, 1985. 55(3): p. 836–9.

59. Kolaskar, A.S. and P.C. Tongaonkar, A semi-empirical method for prediction of antigenic determinants on protein antigens. FEBS Lett, 1990. 276(1-2): p. 172–4.

60. Ponomarenko, J., et al., ElliPro: a new structure-based tool for the prediction of antibody epitopes. BMC Bioinformatics, 2008. 9: p. 514.

61. Andreatta, M. and M. Nielsen, Gapped sequence alignment using artificial neural networks: application to the MHC class I system. Bioinformatics, 2016. 32(4): p. 511–7.

62. Buus, S., et al., Sensitive quantitative predictions of peptide-MHC binding by a ‘Query by Committee’ artificial neural network approach. Tissue Antigens, 2003. 62(5): p. 378–84.

63. Lundegaard, C., et al., NetMHC-3.0: accurate web accessible predictions of human, mouse and monkey MHC class I affinities for peptides of length 8-11. Nucleic Acids Res, 2008. 36(Web Server issue): p. W509–12.

64. Lundegaard, C., O. Lund, and M. Nielsen, Accurate approximation method for prediction of class I MHC affinities for peptides of length 8, 10 and 11 using prediction tools trained on 9mers. Bioinformatics, 2008. 24(11): p. 1397–8.

65. Lundegaard, C., M. Nielsen, and O. Lund, The validity of predicted T-cell epitopes. Trends Biotechnol, 2006. 24(12): p. 537–8.

66. Nielsen, M., et al., Reliable prediction of T-cell epitopes using neural networks with novel sequence representations. Protein Sci, 2003. 12(5): p. 1007–17.

67. Nielsen, M. and O. Lund, NN-align. An artificial neural network-based alignment algorithm for MHC class II peptide binding prediction. BMC Bioinformatics, 2009. 10: p. 296.

68. Bui, H.H., et al., Predicting population coverage of T-cell epitope-based diagnostics and vaccines. BMC Bioinformatics, 2006. 7: p. 153.

69. Chen, J.E., C.C. Huang, and T.E. Ferrin, RRDistMaps: a UCSF Chimera tool for viewing and comparing protein distance maps. Bioinformatics, 2015. 31(9): p. 1484–6.

70. Hertig, S., et al., Multidomain Assembler (MDA) Generates Models of Large Multidomain Proteins. Biophys J, 2015. 108(9): p. 2097–102.

71. Kallberg, M., et al., Template-based protein structure modeling using the RaptorX web server. Nat Protoc, 2012. 7(8): p. 1511–22.

72. Pettersen, E.F., et al., UCSF Chimera--a visualization system for exploratory research and analysis. J Comput Chem, 2004. 25(13): p. 1605–12.

73. Yang, Z., et al., UCSF Chimera, MODELLER, and IMP: an integrated modeling system. J Struct Biol, 2012. 179(3): p. 269–78.

74. Gasteiger, E., et al., ExPASy: The proteomics server for in-depth protein knowledge and analysis. Nucleic acids research, 2003. 31: p. 3784–8.

75. Inc CCG. Molecular Operating Environment (MOE). 2007; Available from: http://www.chemcomp.com.

76. BIOVIA DS. Discovery Studio Visualizer 2.5 ed. San Diego: Dassault Systèmes. 2009; Available from: https://www.3dsbiovia.com/.

77. Shen, Y., et al., Improved PEP-FOLD Approach for Peptide and Miniprotein Structure Prediction. J Chem Theory Comput, 2014. 10(10): p. 4745–58.

78. Thevenet, P., et al., PEP-FOLD: an updated de novo structure prediction server for both linear and disulfide bonded cyclic peptides. Nucleic Acids Res, 2012. 40(Web Server issue): p. W288–93.

79. Pfaller, M.A. and D.J. Diekema, Epidemiology of invasive candidiasis: a persistent public health problem. Clin Microbiol Rev, 2007. 20(1): p. 133–63.

80. de Klerk, N., et al., Fructose-bisphosphate aldolase and pyruvate kinase, two novel immunogens in Madurella mycetomatis. Med Mycol, 2012. 50(2): p. 143–51.

81. Elhag Mustafa, M., et al., Immunoinformatics Prediction of Epitope Based Peptide Vaccine Against Listeria Monocytogenes Fructose Bisphosphate Aldolase Protein. 2019.

82. Elhag Mustafa, M., et al., Design of Epitope Based Peptide Vaccine Against Pseudomonas Aeruginosa Fructose Bisphosphate Aldolase Protein using Immunoinformatics. 2019.

83. Elhag Mustafa, M., et al., Immunoinformatics Prediction of Epitope Based Peptide Vaccine Against Schistosoma Mansoni Fructose Bisphosphate Aldolase Protein. 2019.

84. Mohammed, A.A., et al., Epitope-Based Peptide Vaccine Against Fructose-Bisphosphate Aldolase of Madurella mycetomatis Using Immunoinformatics Approaches. Bioinform Biol Insights, 2018. 12: p. 1177932218809703.

85. Testa, J.S. and R. Philip, Role of T-cell epitope-based vaccine in prophylactic and therapeutic applications. Future Virol, 2012. 7(11): p. 1077–1088.

86. Ab, I., et al., Immunoinformatics Predication and Modelling of a Cocktail of B-and T-cells Epitopes from Envelope Glycoprotein and Nucleocapsid Proteins of Sin Nombre Virus. Immunome Res 2017. 13: p. 141.

